# Mutant TP53 switches therapeutic vulnerability during gastric cancer progression within Interleukin-6 family cytokines

**DOI:** 10.1101/2024.04.22.590499

**Authors:** Anne Huber, Amr H. Allam, Christine Dijkstra, Stefan Thiem, Jennifer Huynh, Ashleigh R. Poh, Joshua Konecnik, Saumya P. Jacob, Rita Busuttil, Yang Liao, David Chisanga, Wei Shi, Mariah G. Alorro, Stephen Forrow, Daniele V.F. Tauriello, Eduard Batlle, Alex Boussioutas, David S. Williams, Michael Buchert, Matthias Ernst, Moritz F. Eissmann

## Abstract

Although aberrant activation of the KRAS and PI3K pathway alongside TP53 mutations account for frequent aberrations in human gastric cancers, neither the sequence nor the individual contributions of these mutations have been clarified. Here, we establish an allelic series of mice to afford conditional expression in glandular epithelium of *Kras^G12D^*; *Pik3ca^H1047R^* or *Trp53^R172H^* and/or ablation of *Pten* or *Trp53*. We find that *Kras^G12D^*;*Pik3ca^H1047R^*is sufficient to induce adenomas, and that lesions progress to carcinoma when also harboring *Pten*-deletions. Additional challenge with either *Trp53* loss- or gain-of-function alleles further accelerated tumor progression and triggered metastatic disease. While tumor-intrinsic STAT3 signaling in response to gp130 family cytokines remained as a gatekeeper for all stages of tumor development, metastatic progression required a mutant *Trp53*-induced interleukin (IL)-11 to IL-6 dependency switch. Consistent with poorer survival of patients with high IL6 expression, we identify IL6/STAT3 signaling as a therapeutic vulnerability for TP53-mutant gastric cancer.

---

Gastric cancer (GC) accounts for the fifth most diaggresnosed and the third most common cause of cancer related death worldwide ^1^. Because a majority of GC is first diagnosed when patients present with distal metastasis, the overall 1-year survival for GC patients remains below 25% ^2^. With potentially curative surgery often not possible for metastatic GC patients, current therapies have only limited life prolonging effects. In part this is due to the heterogeneity of GC at the adenocarcinoma stage, where many somatic mutations drive mutagenesis and disease progression.

Large whole genome sequencing studies suggested specific gene mutation frequencies and allowed definition of molecular based GC subtypes ^3-5^, consistent with the histological Lauren classification system categorizing GC in diffuse, intestinal, and mixed subtypes. While alterations in the gene encoding tumor suppression protein (*TP53*) accounting for the most common mutations across all molecular subtypes, the chromosomally instable (CIN) subtype contributes half of all GC. In addition to *TP53* mutations, the CIN subtype is often associated with amplification of receptor tyrosine kinases and associated Ras signaling, and to a lesser extent with PIK3CA pathway alterations. Unlike the most common forms of colon cancer, where the availability of early stage, non-invasive lesion enabled the establishment of the sequence of genetic events underpinning disease, the sequence of mutations resulting in invasive GC is less clear. In particular, the requirement for activating mutations in the canonical WNT signaling pathway, including loss of function mutations in its primary negative regulator, the APC tumor suppressor gene, during the ontogeny of GC, remains controversial. While *APC*-mutation associated with excessive activation of canonical WNT signaling accounts for the initiating event in a large majority of the most common sporadic forms of colon cancer, they occur in less than 10% of GCs. However, gene signatures associated with excessive activation of the WNT/b-catenin signaling pathway are associated with 80% of GC ^6^. On the other hand, mutations in TP53 account for relatively late events in most epithelial malignancies and are often placed at the stage when adenocarcinomas acquire aggressive metastatic characteristics.

Mutations in the *TP53* protein have been proposed to promote tumor progression as a consequence of three potentially overlapping outcomes. Besides the complete loss-of-function, expression of mutant proteins occurs. They predominantly arise from missense mutations in hotspots located in the DNA binding domain of the protein thereby impacting on the formation of the transcriptionally active TP53 tetramer. Accordingly, many mutant forms of TP53 are likely to exert a dominant-negative function on wildtype TP53, while others, including the most prevalent *R175H* missense mutation in human TP53 (equivalent to *Trp53^R172H^* in mice ^7,8^) may result in gain-of-function consequences as they exacerbate invasion and metastasis ^9^. However, with either type of mutation occurring in one allele, the remaining wild-type (wt) allele is frequently lost through deletion of large chromosomal fragments resulting in loss-of-heterozygosity (LOH) and associated functional balancing of the remaining wild-type TP53 protein ^10^.

Due to its role as the guardian of the genome, it has been suggested that TP53 mutations may result in novel vulnerabilities of cancer cells that can be exploited therapeutically, including the appearance of novel tumor neo-antigens ^11-13^. On the other hand, TP53 mutation-dependent transcriptional changes within tumor cells may also lead to novel “addictions” to non-mutated signaling pathways. Notably, wild-type TP53 is a transcriptional suppressor of interleukin (IL)6 ^14,15^, while the presence of either gain- or loss-of-function TP53 mutations increased IL6 expression and activation of the associated signaling pathway, comprising the shared GP130 receptor subunit and STAT3 as the transcriptional signaling node ^16-18^. However, early adenoma stages are already fueled by aberrant STAT3 activity as a result of oversupply of inflammatory cytokines, which is often observed even in the absence of overt gastritis ^19^.

Indeed, the GP130 family cytokine IL11 rather than IL6 becomes rate-limiting for the growth of intestinal-type gastric cancer, at least during adenomatous stages in autochthonous mouse models^20-23^. On the other hand, elevated STAT3 in the stromal cells of the host confers an immune-suppressed tumor microenvironment with specific roles identified for IL6 and IL11. While the former cytokine helps setting up a premetastatic niches ^24^, signaling from the latter suppresses the activity of CD4 cells and antagonizes the host’s anti-tumor immune response ^25^. Meanwhile, high STAT3 activity in human patients correlates with GC progression, metastasis, and poor patient survival ^26^.

Here we provide a novel allelic series of autochthonous models for metastasizing intestinal-type GC that occurs in the absence of activating mutations in the WNT pathway. We identify a critical role for TP53 mutations, irrespective of their functional consequences, for the transition between non-invasive adenomas to metastasizing carcinomas. This functionally correlates with a switch from IL11 to IL6 dependency. Surprisingly, the requirement for IL6 remains intrinsic to cancer cells and transplantable via the corresponding tumor organoids thereby highlighting opportunities to discover novel therapeutic vulnerabilities over and above the addiction to IL6 signaling identified here.

## Results

### WNT-signaling independent *Kras*, *Pik3ca* and *Trp53* mutation-driven model of invasive and metastatic stomach adenocarcinoma

Because mutations in multiple common oncogenes and tumor suppressors underpin aberrant activity of signaling pathways that contribute to GC progression ^3-5,27^, we re-analyzed the TCGA stomach adenocarcinoma dataset for the 10 most frequently involved pathways ^27^ (Supplementary Fig. 1A). We identified the cell cycle as being the most frequent subject of mutations, followed by alterations to the Receptor Tyrosine Kinase (RTK)/RAS, PI3K and TP53 pathways, where simultaneous mutations within the latter three occurs in 15.1% of stomach adenocarcinoma (STAD) patients (Supplementary Fig. 1B). Since aberrant pathway activity can occur independently of mutations in corresponding genes, we confirmed that 48.3% of STAD patients simultaneously display elevated transcriptional activation in the RTK/KRAS and the PI3K pathway, and 23% in RTK/KRAS, PI3K and TP53 pathways (Supplementary Fig. 1C, D). We also noted that 37.3% of all STAD patients showed mutations in the canonical WNT signaling pathway (Supplementary Fig. 1A).

To establish corresponding mouse models, we exploited our BAC-transgenic and tamoxifen-inducible *Tff1:CreERT2* driver strain ^28^ to conditionally induce various combinations of latent activatable alleles to encode KRAS^G12D^, PIK3ca^H1047R^ or TP53^R172H^ mutant proteins, alongside the deletion of PTEN through induction of the *Pten*^del^ allele. While we previously described that gastric epithelial specific expression of *Kras^G12D^* is sufficient to trigger gastric adenoma formation ^28^, we only detected adenomas when concurrently mutating *Pi3kca* and *Pten* with the *Tff1^CreERT2^* driver stain, but not when either gene was mutated individually (Supplementary Fig. 1E-H). Meanwhile, 19% of tamoxifen-induced compound mutant *Tff1^CreERT2^*;*Kras^G12D^*;*Pik3ca^H1047R^*(referred to as *KP*) mice developed gastric tumors and 9.5% presented with adenocarcinomas (Fig. 1A). However, further augmenting PI3K pathway activation through heterozygous ablation of *Pten* in triple mutant *Tff1^CreERT2^*;*Kras^G12D^;Pik3ca^H1047R^;Pten^del^* (*KPP*) mice increased the overall frequency of gastric tumors to 81%, and over half had progressed to carcinomas. Interestingly, neither lesions in *KP* nor *KPP* mice progressed to metastatic stages (Fig. 1A, B). Indeed, despite the larger size of *KPP* tumors when compared to their *KP* counterparts, both types of tumors were characterized by elongated pits alongside enlarged glandular structures associated with accumulation of intraepithelial lymphocytes, while submucosal invasion was more evident in *KPP* tumors (Fig. 1B).

**Fig. 1.**
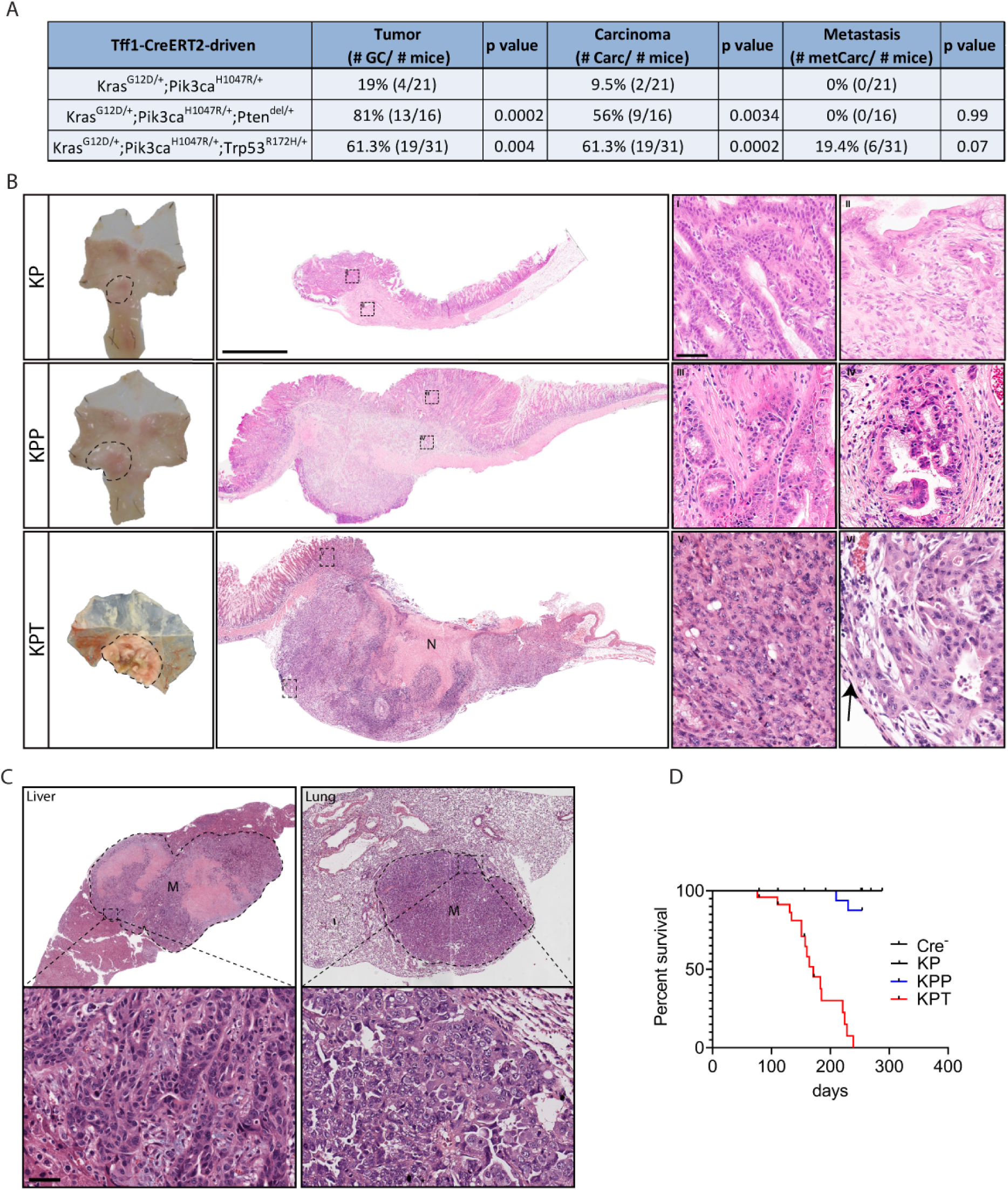
Mutant Kras, Pik3ca and Trp53 drive gastric invasive carcinoma formation. **A** Table summarizing gastric tumor (GC), carcinoma (Carc), and metastatic carcinoma (metCarc) incidences after tamoxifen administration to *Tff1^CreERT2^* positive mice harboring the lox-STOP-lox (LSL)-flanked exons *Kras^LSL-G12D/+^;Pik3ca^LSL-H1047R/+^*(*KP*), *Kras^LSL-G12D/+^;Pik3ca^LSL-H1047R/+^;Pten^flox/+^*(*KPP*) or *Kras^LSL-G12D/+^;Pik3ca^LSL-H1047R/+^;Trp53^LSL-R172H/+^* (*KPT*). P values of Fisher’s exact test are shown. **B** Representative images of whole mount stomachs (left) and microscopic images of H&E stained stomach cross sections (scale bar = 2mm) containing lesion from mice with genotypes as in **A**. I-VI show higher power images of mucosal and submucosal (=invasive) parts of the tumors (scale bar = 50µm). N depicts necrotic tumor tissue and the arrow (in VI) points at tumors cells invading stomach serosa. **C** Representative H&E stained liver (left) and lung (right) sections containing metastasis in KPT mice (scale bar = 50µm). **D** Kaplan-Meier survival analysis of mice with genotypes: Tff1^CreERT2^ negative (Cre^-^) and Tff1^CreERT2^ positive mice harboring the mutant allele combinations: *Kras^LSL-G12D/+^;Pik3ca^LSL-H1047R/+^* (*KP*), *Kras^LSL-G12D/+^;Pik3ca^LSL-H1047R/+^;Pten^flox/+^*(*KPP*) or *Kras^LSL-G12D/+^;Pik3ca^LSL-H1047R/+^;Trp53^LSL-R172H/+^* (*KPT*). Log-rank (Mantel-Cox) test for KPP vs KPT results are HR=0.063 (0.024 – 0.168) and p<0.0001. N = 16, 16, 16 and 24 respectively.

Next, we challenged *KP* mice with additional TP53 mutations as a later stage event in STAD patients associated with tumor progression. Indeed, simultaneous gene (in-)activation in the resulting *Tff1^CreERT2^*;*Kras^G12D^*;*Pik3ca^H1047R^*;*Trp53^R172H^*(*KPT*) mice, transformed the lesions observed in *KPP* mice to adenocarcinomas of the intestinal type. *KPT* tumors were classified as poorly differentiated tubular-type adenocarcinomas and showed extensive lymphatic and submucosal invasion, which was frequently associated with tissue necrosis (Fig. 1A, B, and Supplementary Fig. 1I, Supplementary Table 1). One third of moribund tumor-bearing *KPT* mice presented with liver and lung metastases (Fig. 1A, C), which correlated with reduced overall survival when compared to mice from *KPP* and *KP* cohorts (Fig. 1D).

Given the high prevalence of mutations in the canonical WNT signaling pathway in human GC, we excluded contribution by secondary serendipitous mutations in the pathway to tumor formation in *KPP* and *KPT* mice using the absence of nuclear β-catenin as a surrogate marker for activation of the canonical WNT pathway (Supplementary Fig. 2A). Likewise, we could not detect activation of the canonical WNT-target and stem cell genes *Lgr5* and *Sox9* but noted increased expression of the more promiscuous genes *CD44*, *Ccnd1* and *Myc* in *KPT* tumors (Supplementary Fig. 2B), which have also been identified as targets for STAT3.

### Loss-of-function and gain-of-function *Trp53* mutations can drive aggressive metastatic disease

TP53 mutations can be classified to either have loss-of-function or possible gain-of-function consequences ^9^. Although the direct relationship between specific amino acid substitutions in *Tpr53* and functional outcome remains controversial, LOH of the remaining wild-type allele is a frequent consequence. Prior to Cre-mediated recombination, the lox-STOP-lox cassette within intron 1 of the targeted *Trp53^R172H^* locus blocks the expression from that allele, thereby effectively constitutes a “Loss-of-Expression” allele (subsequently referred to as *Trp53^LoE^*) ^8^. However, upon Cre-mediated recombination this allele is reconstituted to contain the R172H substitution, which corresponds to the R175H hotspot gain-of-function mutations found in human cancer patients (subsequently referred to as *Trp53^GoF^*) (Supplementary Fig. 3). Due to the incomplete activity of Cre-recombinase and the aforementioned LOH observations, we clarified the *Trp53* status in gastric carcinomas and tumor organoids established from *KPT* mice (Fig. 2A). We detected all four possible variants of *Trp53* allele combinations, without any apparent preference for one status over another (Fig. 2B). Strikingly, *Trp53^LoE/-^* or *Trp53^GoF/-^* tumor-bearing mice, showed reduced survival when compared to littermates with *Trp53^LoE/Wt^* or *Trp53^GoF/Wt^*tumors that still harbored a wild-type allele (Fig. 2C). Meanwhile, primary tumors of mice with synchronous metastases returned all but the *Trp53^GoF/Wt^* allele combination in their primary lesions (Fig. 2D).

**Fig. 2.**
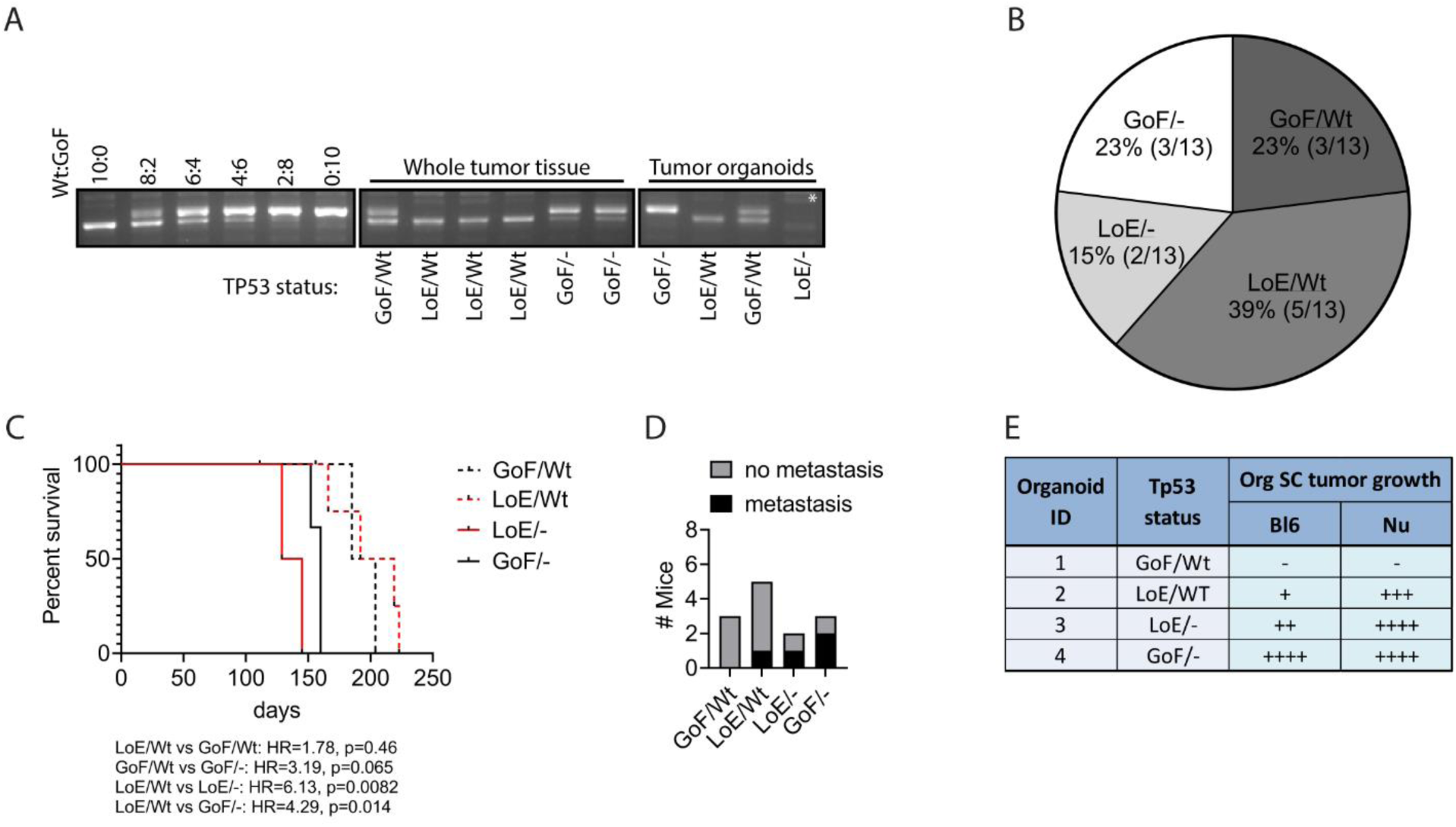
Tp53 mutation status in KPT tumors. **A** Image of *Trp53* status PCR assay performed on genomic DNA of whole tumor or organoid lysates from *Tff1^CreERT2/+^;Kras^LSL-G12D/+^;Pik3ca^LSL-H1047R/+^*;*Trp53^LSL-R172H/+^*(*KPT*) mice. GoF = Gain-of-Function = recombined *Trp53^LSL-R172H^*allele, LoE = Loss-of-Expression = non –recombined *Trp53^LSL-R172H^*allele leads to no *Trp53* being expressed, Wt = wild type *Trp53*, - = the wild type *Trp53* allele being genetically lost. Asterisk (*) indicates that secondary genomic DNA PCR was used to confirm presence of the non-recombined *Trp53^LSL-R172H^* allele. **B** Frequency of *Trp53* status assessed by PCR assay (shown in A) in tumors from *KPT* mice. **C** Kaplan-Meier survival analysis of *KPT* mice based on tumor *Trp53* status. Hazard ratios (HR) and p values of Log-rank (Mantel-Cox) test are shown. N = 3, 5, 2 and 3 respectively. **D** Number of KPT mice presenting with or without metastasis based on primary tumor *Trp53* status. **E** Table depicting the tumor allograft growth potential of gastric cancer organoids derived from *KPT* mice with indicated Trp53 status in C57/BL6 wild type and BALB/c Nu/Nu host mice (n=4 host mice per organoids per background). - = no tumor growth within 80days, + = initial tumor ≤ 50mm^3^ form but do not progress within 80 days, ++ = tumors grow ≥1000mm^3^ within 80 days, +++ = tumors grow ≥1000mm^3^ within 55 days, ++++ = tumors grow ≥1000mm^3^ within 30 days.

In order to address whether these allele combinations functionally contributed to the growth characteristics of primary *KPT* tumors, we established organoid cultures from independent tumors to cover all four allele combinations and investigated their growth characteristics when grown as allograft tumors in either immune competent C57BL/6J wild-type mice, or in immune-deficient BALB/c Nude mice (Fig. 2E). Strikingly, allografts consistently grew quickest when established from tumor organoids that lacked a *Trp53^Wt^*allele, and slowest when the altered allele was still balanced with its wild-type counterpart irrespective of the immune status of the host. We surmise from these observations that both loss-of-expression and *Trp53^R172H^*-encoded putative gain-of-function mutations promote tumor growth and progression and that this is further exaggerated through the loss of the *Trp53^Wt^* allele.

### TP53 dependent switch for mandatory STAT3 activity in tumor cells from IL11 to IL6

We and others have previously demonstrated that STAT3 signaling in response to IL6 family cytokines provides a rate-limiting signal for gastrointestinal tumors that arise from *bona fide* oncogenic driver mutations, including in *APC*, *KRAS* or other genes ^20-23,29^. Indeed, using the nuclear presence of the phosphorylated STAT3 (p-STAT3) isoform as a surrogate, we identified active STAT3 signaling in epithelial cells localized in the mucosa as well as the submucosal invasive fronts across gastric tumors of *KPP* and *KPT* mice (Fig. 3A). To functionally evaluate this observation, we performed subcutaneous allografts with organoids derived from the invasive fronts of adenocarcinomas from *KPT* mice (Fig. 3B). Treatment with the STAT3-inhibitor BBI608 ^30^ significantly reduced tumor size, suggesting that the growth of *KPT* mutant tumors is fueled by STAT3 signaling (Fig. 3C). We genetically confirmed that the effect of systemic BBI608 administration was at least in part due to tumor cell-intrinsic STAT3 signaling, because CRISPR-Cas9-mediated STAT3 deletion in *KPT* organoids reduced their growth when established as allografts in *Stat3^Wt^* hosts (Fig. 3D, E). The *in vitro* growth characteristics between STAT3-proficient and STAT3-deficient *KPT* organoids remained indistinguishable in the absence of exogenously added gp130 family cytokines (Supplementary Fig. 4A).

**Fig. 3.**
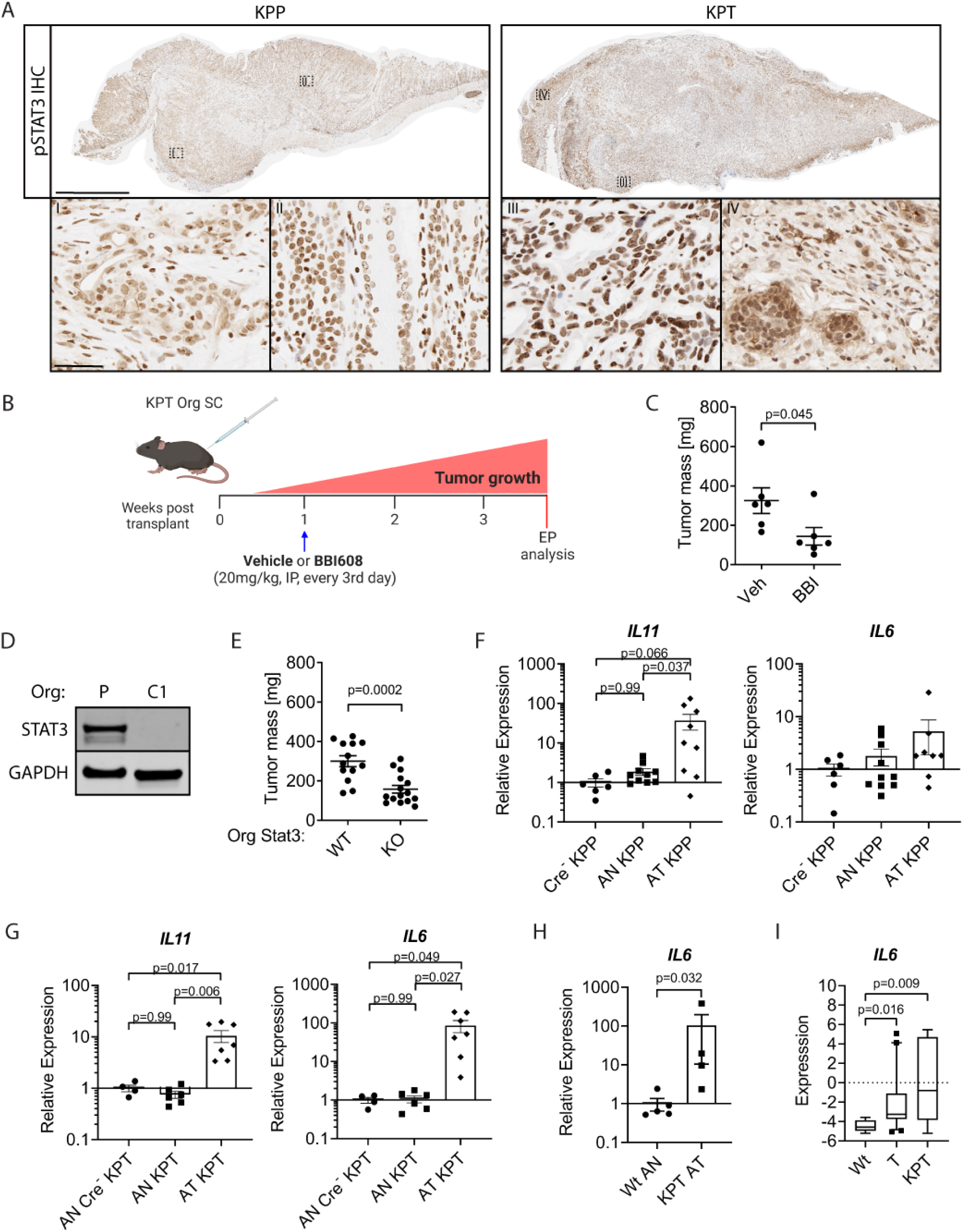
Stage-specific ligand switch for mandatory Stat3 activity in tumor cells. **A** Representative photograph of pSTAT3 immunohistochemistry staining of a *Tff1^CreRT2^; Kras^LSL-G12D/+^; Pik3caLSL-H1047R/+; Ptenflox/+* (*KPP*; left) and a *Tff1CreRT2; KrasLSL-G12D/+; Pik3caLSL-H1047R/+; Trp53LSL-R172H/+* (*KPT*; right) mouse stomachs (scale bar = 2mm) with magnification of the invasive tumor front (I and III) and the mucosal tumor core (II and IV) (scale bar = 50µm). **B** Schematic of *KPT* mutant GC organoid (Org) subcutaneous (SC) transplantation with indicated treatment cohorts, EP-endpoint. Created with Biorender.com. **C** Tumor mass at endpoint of *KPT* Org SC allograft experiment as outlined in **B** for vehicle (Veh) and BBI608 (BBI) treated animals. N = 6 and 6 (experiment was performed once). **D** Immunoblotting for STAT3 and actin protein on organoid lysates from parental (P) and CRISPR-Cas9 Stat3^KO^ clone 1 (C1). **E** Tumor mass at endpoint of Org SC allograft experiment, where either Stat3^WT^ (Wt) or Stat3^KO^ (KO) *KPT* GC organoids were transplanted. N = 13 and 15 (pooled data from two independent experiments). **F,G** qPCR gene expression analysis for *Il11* and *Il6* in whole tissue lysates of antrum normal tissue (AN) and antrum tumor (AT) of indicated genotypes. Expression data is presented relative to mean of Cre^-^ KPP (**F**) or AN Cre^-^ KPT (**G**) data points. P values shown are fromo-way ANOVA + Turkey’s multiple comparison testing; for the IL6 graph (in **F**) all p>0.3. N = 6, 10 and 9 (**F** left) and n = 6, 10 and 8 (**F** right). N = 4, 6 and 7 (**G**, both graphs). **H** qPCR-determined expression levels of *Il6* in organoids derived from wild type antrum stomach (Wt) and stomach tumors of KPT mutant mice. Expression data is presented relative to Wt AN values. P-values of Mann Whitney test are shown. N = 5 and 4. **I** Expression levels of *IL6* in human gastric cancer cell lines grouped into TP53^Wt^ (Wt), TP53 (T) mutant or <u>K</u>RAS;<u>P</u>I3K;<u>T</u>P53 (*KPT*) mutant activation signature positive. RNA sequencing data was downloaded from the Broad Institutes Cancer Cell Line Encyclopedia. Data is shown as Box & Whiskers plots (10-90 percentile), n=8, 21 and 7 respectively. Kruskal-Wallis and Dunn’s multiple comparison’s test were performed. Data represents mean±SEM (**C, E-I**). Each symbol represents a biological replicate, specifically one mouse (**C**, **E**-**G**), independent organoid culture (**H**) or human GC cell line (**I**). Two-sided Student’s *t*-test p values are shown (**C, E**).

We next aimed to identify the *in vivo* contribution of individual GP130 cytokines on the growth of *KPP* and *KPT* tumors, respectively. *IL11* was the most prominently expressed GP130-cytokine in *KPP* tumors when compared to adjacent non-tumor epithelium, while IL6 expression remained comparable between these two compartments (Fig. 3F). By contrast, we also detected elevated *IL6* expression in *KPT* tumors (Fig. 3G). Importantly, *IL6* expression was elevated in *KPT* organoids (Fig. 3H), reminiscent of the increased IL6 expression in patient derived human TP53 mutant and *KPT* gastric cancer cell lines (Fig. 3I). Indeed, co-culture of bone marrow-derived macrophages (BMDM) with *KPT* organoids increased *IL6* expression as long as the *KPT* organoids harbored a mutated *Trp53* allele (Supplementary Fig. 4B). Surprisingly, the presence of the *Trp53* mutant alleles was specifically associated with elevated *IL6* but not *IL11* expression in *KPT* organoids, human GC cell lines and in BMDM-organoid co-cultures (Supplementary Fig. 4C-E). These results suggest that the elevated IL6 associated with TP53 mutant tumors may be derived from the tumor cells as well as cells of the microenvironment, such as macrophages.

To confirm whether IL11-mediated Stat3 signaling is involved in the formation and progression of the gastric lesions, we generated *KPP*;*IL11ra^+/-^*mice, because we had previously shown that monoallelic reduction of the IL-11Ra receptor subunit impaired IL11-dependent formation of a signaling-competent IL11:IL11Ra:gp130 receptor complexes ^21^. Indeed, heterozygous *IL11ra* ablation reduced tumor size and overall frequency compared to tumors from *KPP*;*IL11ra^+/+^* litter mates, as well as preventing progression of tumors in *KPP*;*IL11ra^+/-^* mice to carcinoma stages (Fig. 4A-D). In stark contrast, the incidence of carcinoma, their depth of invasion and metastatic capacity remained comparable between *Trp53*-mutant *KPT*;*IL11ra^+/-^* and *KPT*;*IL11ra^+/+^*littermates (Fig. 4E and Supplementary Fig. 5A). Consistent with this observation, pSTAT3 protein levels and nuclear staining remained comparable between tumors of *KPT;IL11ra^+/-^*and *KPT*;*IL11ra^+/+^* mice, while pSTAT3 was decreased in tumors from *KPP;IL11ra^+/-^* compared to those from *KPP*;*IL11ra^+/+^*mice (Supplementary Fig. 5B-E). Since we detected increased *IL6* expression in tumors of *KPT* mice, we also assessed the causal consequences of this correlation and found that *KPT* organoid tumor allografts grew slower in IL6-deficient hosts (Fig. 4F). While therapeutic administration of neutralizing IL6 antibodies to wild-type hosts with established *KPT* tumor allografts reduced their growth (Fig. 4G), this was not observed when *KPT* organoids were implanted in either IL11 ligand- or IL11Ra receptor-deficient hosts (Fig. 3H, I). Collectively, our data established tumor intrinsic STAT3 signaling as a rate-limiting gate-keeper function for *KPT* tumors and suggests a switch from IL11-dependency by *Trp53^Wt^* tumors to IL6-dependency once *Trp53* has acquired oncogenic mutations.

**Fig. 4.**
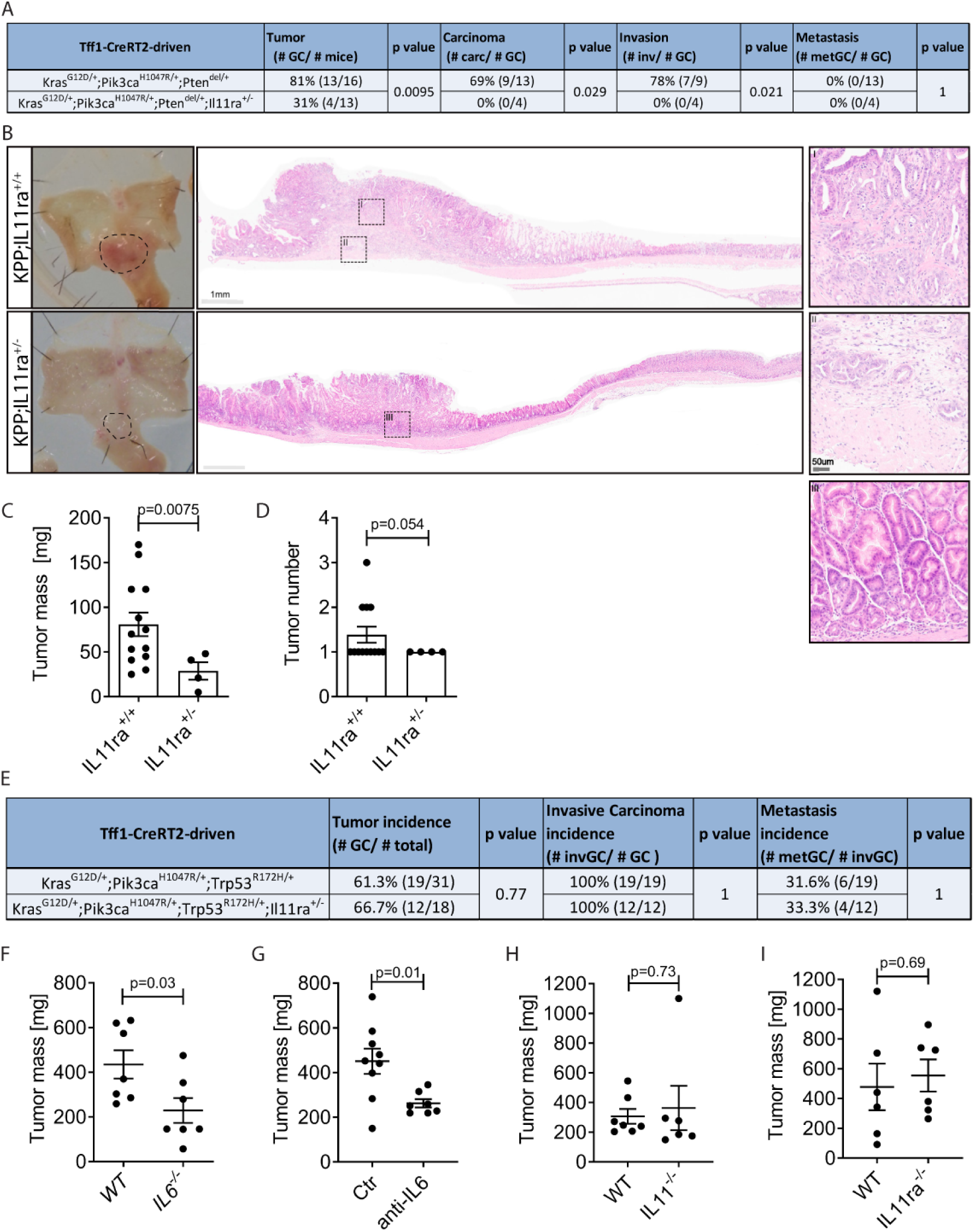
Functional stage-specific reliance on IL6 family cytokines. **A** Table summarizing gastric tumor (GC), carcinoma (carc), cancer invasion (inv) and metastatic GC (metGC) incidences after tamoxifen administration to *Tff1^CreRT2^* positive mice either harboring *Kras^LSL-^G12D/+; Pik3caLSL-H1047R/+*;*Ptenflox/+* (*KPP*) or *KrasLSL-G12D/+; Pik3caLSL-H1047R/+*;*Ptenflox/+;Il11ra+/- (KPP;IL11ra+/-*). *KPP* data are also shown in Fig. 1A. **B** Representative HE staining of whole mount stomachs and microscopic images of stomach lesion and of mice with genotypes as in **A**. Scale bars are as indicated. **C,D** Gastric tumor burden (**C**) and tumor number (**D**) analysis is shown from tumor-bearing mice (*Tff1^CreERT2/+^; Kras^LSL-G12D/+^; Pik3ca^LSL-H1047R/+^*;*Pten^flox/+^*) either *IL11ra^+/+^* or *IL11ra^+/-^* at 250 days post mutant allele induction. N = 13, 4 (**C**, **D**). **E** Table summarizing gastric tumor (GC), carcinoma (carc), cancer invasion (inv) and metastatic GC (metGC) incidences after tamoxifen administration to *Tff1^CreRT2^* positive mice either harboring *Kras^LSL-^ G12D/+; Pik3caLSL-H1047R/+*;*Trp53LSL-R172H/+* (*KPT*) or *KrasLSL-G12D/+; Pik3caLSL-H1047R/+*;*Trp53R172H/+;Il11ra+/-* (*KPT;IL11ra^+/-^*). *KPT* data are also shown in Fig. 1A. **F-I** Tumor mass at endpoint analysis of organoid subcutaneous allograft experiment, where *KPT* GC organoids were transplanted into either WT or IL6^-/-^ mice (**F**), WT mice and were treated when tumors reached 100 mm^3^ volume with isotype control antibody (Ctr) or with anti-IL6 neutralizing antibody (anti-IL6) (**G**), WT or IL11^-/-^ (**H**) or WT or IL11ra^-/-^ mice (**I**). Experiments were performed once. N = 7, 7 (**F**), 9, 7 (**G**), 7, 6 (**H**) and 6, 6 (**I**). Data represents mean±SEM (**C, D, F-I**). Each symbol represents a biological replicate, specifically one mouse (**C, D, F-I**). P values of Fisher exact test (**A**, **E**) and two-sided Student’s *t*-test (**C, D, F-I**) are shown.

### Elevated IL6 and STAT3 signaling predicts poor survival in gastric adenocarcinoma patients

In order to translate the causal relationship between elevated STAT3 signaling activity and tumor progression in our murine *KPP* and *KPT* models, we interrogated STAD patient samples for evidence of STAT3 activity (Supplementary Table 2). We used the HALO software for unbiased calls between tumor and non-neoplastic surrounding epithelial/stromal cell compartments across tissue arrays of normal and stomach adenocarcinoma tissues (Supplementary Fig. 6A and Fig. 5A). We detected the strongest pSTAT3 signal in the tumor compartment of intestinal-type gastric cancers biopsies (Fig. 5B). However, when quantifying whole tissue cores that included tumor and stromal compartments, we found comparable pSTAT3 between normal and tumor core biopsies across histological subtypes of GC (Supplementary Fig. 6B).

**Fig. 5.**
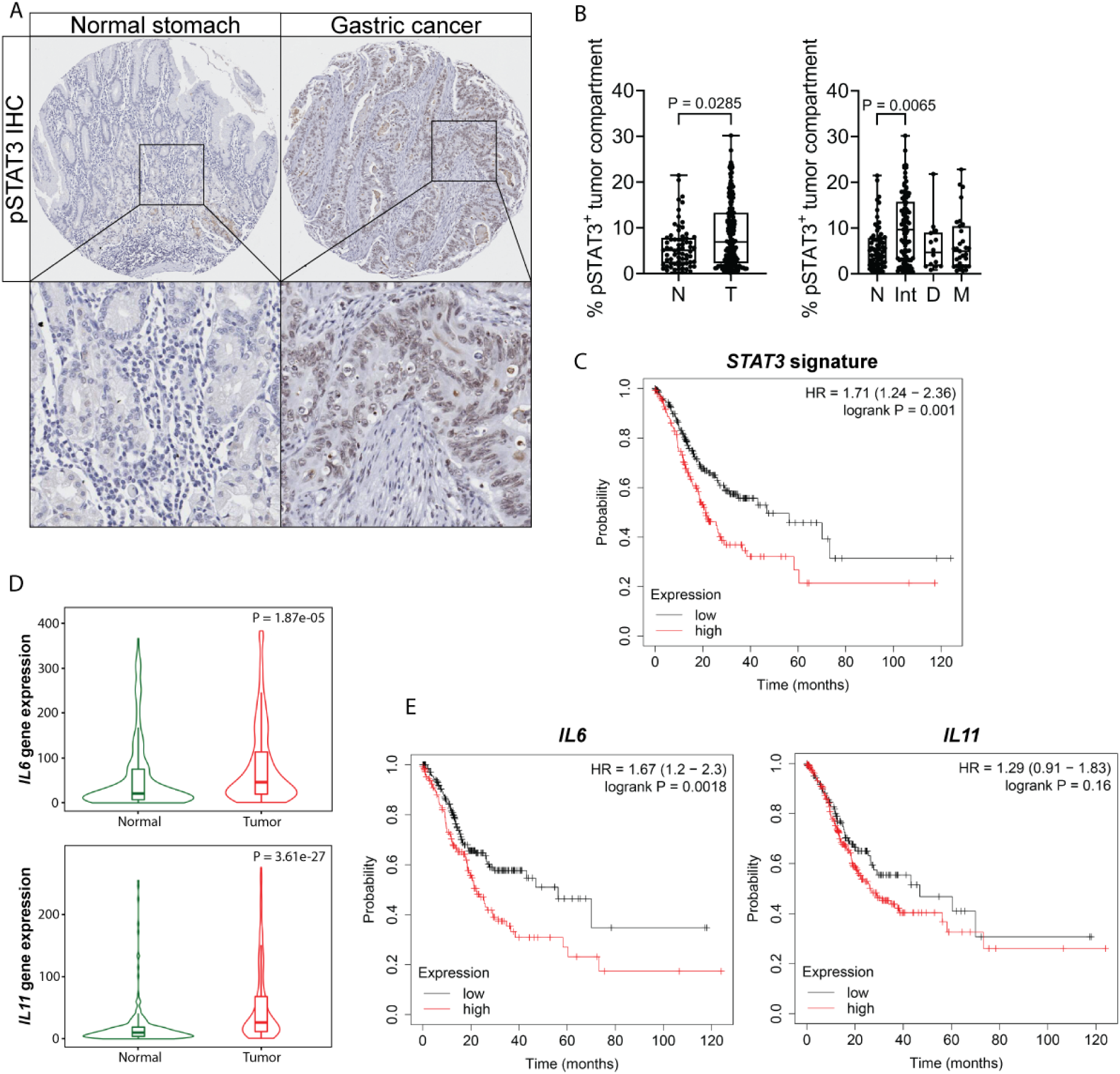
IL6-IL11-Stat3 signaling in gastric cancer patients. **A** Representative image of pSTAT3 immunohistochemical staining on normal stomach and gastric cancer cores of tumor tissue microarray. **B** HALO quantification of pSTAT3 positivity specifically in the tumor compartment comparing normal (N) versus gastric tumor (T) and normal tissues versus gastric cancer Lauren subtypes (right graph; Int=intestinal, D=Diffuse, M=Mixed). Each symbol represents a biological replicate, specifically individual patient samples. N = 68, 154 (left) and 68, 95, 14, 31 (right). **C** Kaplan-Meier survival analysis (overall survival) of STAT3 signaling activation gene signature (STAT3, SOCS3, OSMR, CLD12, PIM3) high versus low in stomach adenocarcinoma (prepared with KM plot). N = 220 (low) and 151 (high). **D** IL6 and IL11 mRNA expression in normal stomach (N) versus gastric tumor (T) tissues. IL6 expression is 1.86 fold and IL11 3 fold increased in the stomach adenocarcinoma (median fold change). P values from Mann-Whitney test (prepared with TNMplot). **E** Kaplan-Meyer Survival analysis (overall survival) IL6 and IL11 high versus low RNA expression in stomach adenocarcinomas (prepared with KM plot). N = 218, 153 (left) and 132, 239 (right). Data represents mean±SEM (**B**). P values of Mann Whitney test (**B left**, **D**), Kruskali-Wallis+Dunnet’s multiple comparisons test (**B** right) and Hazard ratios (HR) and p values of Log-rank (Mantel-Cox) test (**C, E**) are shown.

To independently confirm these observations, we took advantage of our STAT3 gene signature comprised of *bona fide* STAT3-target genes (*STAT3*, *SOCS3*, *CLD12*, *OSMR*, *PIM3*), identified from our STAT3-ChIP-Seq and RNA-Seq data of tumors recovered from *gp130^F/F^* mice stimulated with gp130 cytokines ^31^. When assigning TCGA-patient specimens according to their expression signature into STAT3^Low^ and STAT3^High^ cohorts, Kaplan-Meier survival probability analysis revealed a poorer outcome for the latter (Fig. 5C). In turn, specimen of the STAT3^High^ patient cohort also revealed higher expression of IL6 or IL11 when compared to the STAT3^Low^ cohort (Supplementary Fig. 6C). Furthermore, when compared to normal stomach tissue, gastric cancer specimen displayed elevated expression of both *IL6* and *IL11* (Fig. 5D). However, consistent with the switch from IL11-dependency of mouse *KPP* tumors to IL6-dependency of *KPT* tumors, we found that only high *IL6* expression, but not *IL11* expression correlated with poor patient survival (Fig. 5E). Thus, the gatekeeper role for the IL6-dependent STAT3 signaling cascade for gastric cancer progression in preclinical mouse models shows an equivalent expression correlation in human GC, thereby elevating ligand-specific activation of the GP130/STAT3 signaling cascade as a potential, stage-specific therapeutic target (Supplementary Fig. 7).

## Discussion

Here we create an allelic series of compound mutant mice to enable the inducible formation of autochthonous metastatic tumors that replicate the accumulation of some of the most frequently observed mutations in intestinal-type GC patients. We demonstrate that co-activation of KRAS and PIK3CA/PTEN signaling pathway is sufficient for the growth of gastric tumors to an early adenocarcinoma stage with little invasion and no metastatic progression. Additional mutation in *Trp53* promotes aggressive disease, with all corresponding mice harboring highly invasive adenocarcinomas and frequent distal metastasis, predominantly to the liver. This reflects human disease, where TP53 mutations are found more often in advanced GC involving liver metastasis than in advanced GC without distal metastasis ^32^ and where TP53 mutations are associated with poor prognosis in the microsatellite stable subtype ^4^ and advanced disease ^33^.

The reproducible formation of tumors in *KPP* and *KPT* mice helps to clarify the contribution of aberrant canonical WNT signaling, which has previously been used to underpin a complementing series of GC mouse models based on loss-of-function mutations in the *Apc* tumor suppressor gene ^6^. Interestingly, in the absence of predetermined mutations in the canonical WNT signaling pathway in our *KPP* and *KPT* models, we did not detect nuclear localization of β-catenin, as the surrogate marker for activation of canonical WNT signaling. However, in these tumors we find elevated expression of “promiscuous” WNT target genes, consistent with their transcriptional control through converging RAS/PI3K and STAT3 signaling resulting from the introduced mutations and the IL11/IL6 cytokine production, respectively. Indeed, this is reminiscent of the conversion of the canonical WNT and STAT3 signaling pathways in the context of mutant APC-driven tumor formation in the colonic mucosa ^29^.

The various impacts of genomic alterations in TP53 result from specific mutations causing a broad range of outcomes, including the complete loss of protein function, the generation of proteins with dominant-negative functions, and of proteins with gain-of-function effects ^9^. Although putative gain-of function alterations, including R175H and R248Q, are found in over 20% of the ten most common cancers ^34^, the gain-of-function effect of these missense mutations is likely to only occur in specific cellular contexts^35^. This may explain why recording only the overall TP53 mutation status has proven to be of little prognostic value for GC patients ^36-40^. However, predictive information can be gained when stratifying GC patients according to their molecular subtypes and the nature of their TP53 mutation ^4,33,41-43^, in particular when also including BCL2 Associated X (BAX), Neurexin 1 (NRXN1), Yes Associated Transcriptional Regulator 1 (YAP1) and other additional oncogenic mutations ^38-40^.

Owing to the design of the *Trp53^R172H^* allele and our breeding strategy of *KPT* mice, functional conclusions can be drawn with respect to the presence of the latent “loss-of-expression” or the recombined “gain-of-function” allele, as well as the “balancing” contribution of a *Trp53^Wt^* allele. Notwithstanding our inability to determine whether loss of the *Trp53^Wt^* allele occurs by allelic duplication, we find that the “loss-of-expression” and “gain-of-function” alleles confer similar detrimental effects on the overall tumor burden and survival of *KPT* mice. Meanwhile, loss of a balancing *Trp53^Wt^* allele reduces survival, reminiscent of the LOH being frequently observed at the *TP53* locus in human GC and other cancers ^10,44^ and possibly owing to observations that LOH further accelerates genomic instability and cancer progression ^45^.

Irrespective of the mutation combinations induced in *KPP* or *KPT* mice, we discover a gate-keeper role for tumor cell-intrinsic STAT3 signaling that provides attractive therapeutic targets via interference with the activity of upstream cytokines. Indeed, our results expand on previous observations that excessive STAT3 activity in the gastric epithelium mediated by IL11 triggers the formation of non-invasive adenomas in *gp130^F/F^* mice ^21^. Meanwhile, IL6 and associated STAT3 signaling has recently gained attention as an inflammatory cytokine facilitating metastasis in breast and other solid malignancies possibly by preparing a (pre)-metastatic niche ^24^. TP53 is a transcriptional repressor of IL6 expression ^14,15^, while loss of wild-type TP53 or acquisition of TP53 missense mutations have been shown to increase IL6 and STAT3 signaling ^16-18^. Insight from the switch of cytokine dependency between adenomatous *KPP* tumors expressing wild-type *Trp53* and metastatic *KPT* tumors harboring *Trp53* alterations, suggests significantly different outcomes between tumors exposed *in situ* to IL11 or IL6, despite their shared use of the gp130 co-receptor and associated STAT3 signaling. Indeed, we have previously suggested that IL6, owing to the broader expression of its private receptor subunit IL6Ra compared to IL11Ra, and a phenomenon called IL6 trans-signaling, has a broader spectrum of target cells, including many components of the tumor immune environment that lack expression of the *IL11ra*^21^.

While the exact molecular mechanisms underpinning the TP53 mediated cytokine switch remain not fully understood, RAS-signaling via the AP-1 complex has been shown to transcriptionally activate expression of the *IL11* gene in cancer cells ^46^, while KRAS-mutant cells are poised to coerce cells of the tumor microenvironment into the production of cytokines ^47,48^. On the other hand, loss of TP53 protein is likely to release its transcriptional repressor activity on the IL6 promoter. Meanwhile, the TP53^R172H^ protein has recently been shown to induce expression of the CSF1 cytokine via an interaction with BRD4 on a histone 3 lysine 27 acetylation-rich (H3K27ac) region ^49^, which is also found in a super-enhancer in the *IL6* locus that is susceptible to BRD4-inhibition ^50^. Indeed, epigenetic activation of this super-enhancer, as judged by ATAC sequencing signals, correlated with IL6 expression in esophageal and pancreatic cancer patients. However, TP53^R172H^ may also increase JAK2/STAT3 signaling independent of gp130-cytokines through direct interaction with the gp130-associated phosphatase SHP2 ^17^, reminiscent of the direct binding of the TP53^R248Q^ protein to activated p-STAT3 thereby “short-circuiting” ligand-dependency of the STAT3 signaling cascade during colon cancer progression ^18^.

Although initial clinical trials with the anti-IL6 or IL6Ra monoclonal antibodies Siltuximab and Tocilizumab did not reveal clinical benefits in cancers patients as mono therapies ^51,52^, such trials have neither been conducted in GC patients, nor were patients stratified for their mutational TP53 status. While our study reveals a TP53 mutation-dependent switch from IL11 to IL6 to satisfy the cancer cells’ continuous need for excessive STAT3 activity, our data suggest that the most prominent therapeutic window for interference with IL6 signaling may occur at the stage of initiation/early colonization of the (pre)-premetastatic niche. Further mechanistic insights from the *KPT* model will help to better define recruitment criteria for future clinical trials exploring the beneficial effect of anti-IL6 signaling therapies.

## Methods

### Study approval

Animal experiments were approved and conducted in accordance with all relevant ethical regulations for animal studies including the Australian code for the care and use of animals for scientific purposes. All animal studies were approved by the animal ethics committee of the Ludwig Institute for Cancer Research, the Walter and Eliza Hall Institute of Medical Research, La Trobe University and Austin Health. The conducted research using patient samples was conducted in compliance with all relevant regulations. Collection and usage of human gastric cancer tissues was approved by the Austin Health ethics committee (HREC/15/Austin/359) with a waiver of consent.

### Mice

Mice were co-housed under specific pathogen-free (SPF) conditions and age-and gender-matched littermates were used for experiments. All mouse strains and compound mutants were maintained on C57BL/6J background. Mutant alleles, including those with lox-STOP-lox (LSL)-flanked exons, have been previously described: Tg(Tff1-CreERT2) (hereafter named *Tff1^CreERT2^*)^28^, *Kras^LSL-G12D^*^53^, *Pik3ca^LSL-H1047R^* ^54^, *Pten^flox^* ^55^, *Trp53^LSL-R172H^* mice, 129S-*Trp53tm2Tyj*/J (JAX stock #008652) ^8^, *Apc^fl/fl^* ^56^, *IL6^-/-^* ^57^, *IL11ra^-/-^* ^58^. Compound *Tff1^CreERT2^*;*Apc^fl/fl^* mutant mice are referred to as *Apc^KO^* mice. *Il11^-/-^* mice were generated by EUCOMM/KOMP vector PRPGS00164-A-H10 (containing *Frt*-flanked β-gal reporter and neo selection cassettes, as well as *loxP* sites flanking exons 2–5) targeting in mouse G4 ES cells (C57BL/6Ncr x 129S6/SvEvTac), selected with 200 µg/ml G418. Following screening by long-range PCR and southern blot, 2 clones were injected int C57BL/6J blastocysts. Resultant chimeras (clone 1E9) were mated with wildtype C57BL/6J mice to obtain germline-competent chimeras. These were subsequently mated to *CAGGS-FlpO* transgenic mice and *Sox2-Cre* transgenic mice, to remove *Frt*-flanked selection cassette and *loxP*-flanked *Il11* exons, respectively.

### *In vivo* experiments

#### Cre/lox genetic modified models

Expression of the latent LSL mutant alleles was induced by intraperitoneal (IP) injection of tamoxifen (Sigma-Aldrich, Cat # T5648) in 10% Ethanol, 90% sunflower oil vehicle at 50 mg/kg body weight doses, twice daily for three consecutive days in 6–9-week-old mice that carry the *Tff1^CreERT2^*allele. Upon tamoxifen administration mice were clinically monitored and euthanized at ethical or experimental endpoint (whichever occurred first). Organs of interest were dissected and processes for histology and biochemical and molecular analysis.

#### Subcutaneous GC organoid allograft model

6–8-week-old mice were subcutaneously injected with 900 mechanically disrupted *KPT* GC organoids in 1:1 PBS and RGF BME (R&D Systems, Cat#3533-005-02) vehicle (equivalent of approximately 100,000 cells) into the right flank. An equal number of age-matched male and female host mice were used per experiment. Mice were monitored, and tumors were measured with a caliper (Mitutoyo Tools) three times per week. Tumor volumes were calculated with the formula: (length x (width)^2^)/2. For treatment experiments, drug administration commenced when tumors reached ∼100 mm^3^. BBI608 (Stat3/Stemness inhibitor ^30^, Sellekchem, Cat # S7977) was administered every three days at 20 mg/kg doses in vehicle (5% DMSO, 40% PEG300, 5% Tween80) and anti-IL6 antibody (clone MP5-20FS, BioXCell) was IP injected every three days at 10 mg/kg in PBS vehicle. At experimental endpoint, tumors were dissected and weighed (tumor mass at endpoint).

### Tissue collection

Stomachs, liver, lungs and other organs of interest (for Cre/lox models) and tumors and adjacent tissues (for subcutaneous models) were resected, weighed and then tissue aliquots were snap-frozen for later RNA or protein isolation. Tumor aliquots from the invasive front within the submucosal layers were used to generate GC organoid cultures. The remaining tumor aliquots, tissues and organs were fixed in 10 % neutral buffer formalin and embedded in paraffin blocks for subsequent histological and pathological analysis.

### Histological and pathological analysis

Hematoxylin-Eosin staining of formalin-fixed paraffin-embedded tissues was performed according to “Theory and practice of histological techniques” ^59^.

Pathological assessment was performed by a gastrointestinal pathologist (DW). *KP*, *KPP* and *KPT* mouse tumors were classified using both WHO and Lauren classifications ^60^, and histopathologically assessed in accordance with AJCC cancer staging manual ^61^. Histopathological assessment of *KPT* mouse tumors is summarized in Supplementary Table 1. For the Kaplan-Meier survival analysis (Fig. 1D), *KPT* mice were removed from the analysis when they displayed sarcomas in non-stomach organs or lymphomas in the thymus (n=3) or when they reached ethical endpoint without bearing stomach tumors (n=4).

Osteosarcomas, lymphomas in the thymus and other anatomical sites arise from loss of wild-type TP53 expression in non-stomach body cells ^62^, here through the presence of *Trp53^LoE^* / non-Cre-recombined *Trp53^LSL-172H^*body cells.

### Organoid culture establishment

Mouse gastric tumor organoids were established from the invasive front of intestinal adenocarcinoma bearing *KPT* mice and maintained as previously described ^31,63^.

### CRISPR-Cas9 knockout organoid generation

Stat3^KO^ organoids were established using Alt-R^TM^ CRISPR-Cas9 system (Integrated DNA Technologies), with crisprRNA (ACGATCCGGGCAATTTCCAT) and ATTO^TM^550-labeled tracrRNA. For detailed protocol see Huber *et al* ^64^.

### Organoid- BMDM co-culture experiment

Bone-marrow derived macrophages (BMDM) were established from C57BL/6 wild type mice as previously described ^31^. 1.5×10^5^ day 6 BMDM cells were mixed and seeded together with 300 dissociated GC organoids in 100 µl RGF BME domes in 24-well plate wells and incubated for 72 h in 100 µl IntestiCult^TM^ organoid growth medium (STEMCELL Technologies) containing 20% L929 conditioned medium and 20 ng/ml IL4. At endpoint, growth media were removed, and lysis buffer added to assay wells followed by subsequent RNA isolation and qRT-PCR expression analysis.

### RNA extraction and qRT-PCR analysis

Total RNA from snap frozen tissue was extracted using Trizol^®^Reagent (Life Technologies, Cat# 15596026), while RNA from organoids was extracted using the RNeasy Plus Micro Kit (QIAGEN, Cat# 74034). cDNA was prepared from 2 µg RNA using the High-capacity cDNA Reverse Transcription kit (Applied Biosystems, Cat# 4368813) according to the manufacturer’s protocol.

Quantitative RT-PCR analyses were performed in technical triplicates with SensiMix SYBR kit (Bioline, Cat# QT605–20) using the ViiA™ 7 Real Time PCR System (Life Technologies). Primer sequences used are in Supplementary Table 3.

### Human GC Tissue microarray

Tissue microarrays (TMA) were prepared from gastric (n=193), gastro-esophageal junction (n=66) adenocarcinomas and normal mucosa (n=80; from GC blocks near surgical resection margins, uninvolved by tumor) diagnosed at Austin Health between 2001 and 2014, for whom clinical, treatment and follow-up data had been retrospectively collected with human ethics approval. TMAs were produced from representative FFPE blocks of tumor, sampling 3 x 1 mm cores per patient. Pathology of each core of the TMA was confirmed post generation by a gastrointestinal pathologist (DW). Clinical and pathological characteristics of the patient cohort stratified for P-STAT3 staining positivity are summarized in Supplementary Table 2.

### Immunohistochemical analysis

Paraffin-embedded formalin fixed 4 µm tissue section on charged microscopy glass slides were dewaxed and rehydrated. Antigen retrieval was performed in citrate buffer in a microwave pressure cooker (pH 6 for 15 min). Sections were blocked in 10 % (v/v) normal goat serum for 1 h at 20-25°C in a humidified chamber, incubated overnight at 4°C with primary antibodies and for 1h at room temperature with secondary antibodies. Visualisation was achieved using 3,3-Diaminobenzine (DAB, DAKO). Primary antibodies used were: anti-pSTAT3 at 1.33µg/ml (Tyr705, Cell Signalling, Cat # 9131), anti-β-catenin at 1.25 µg/ml (BD Biosciences, Cat # 610153) and secondary antibodies were: anti-rabbit-biotinylated at 0.75 µg/ml (Vector labs, BA-100, in conjunction with VECTASTAIN ABC kit), anti-mouse-HRP at 5 µg/ml (Dako, Cat # P0447).

For quantification of tissue sections, sections were scanned (Aperio AT2 Leica Scanner) and analysed with Image Scope software (Leica Biosystems, version 12.4).

For the human GC TMAs, HALO software (Indica Labs, version 3.5) was used to distinguish tumour/epithelial and stromal cell compartments (Random Forest Tissue Classifier), as well as to quantify positive staining of cells (Area Quantification Algorithm). Tissue cores that had less than 20% tumor or epithelial cells were removed from tumor/epithelial compartment analysis. Cores with strong non-specific background P-STAT3 staining (predominantly in gastric gland lumen) or cores with greater 40% tissue loss were excluded from the analysis.

### Protein extraction and Immunoblot analysis

Protein lysates from snap-frozen tissue were prepared using the TissueLyser II (Qiagen) and RIPA lysis buffer (Sigma). Cultured organoids were directly lysed by adding RIPA lysis buffer to the organoid domes and mechanical disruption by 20-time pipetting. Immunoblotting was performed as previously described ^31^. Antibodies used were: primary: pSTAT3 (Tyr705, Cell Signalling #9131), STAT3 (Cell Signaling Technology #4904) and GAPDH (Merck #G9545); secondary antibodies: fluorescent-conjugated (LI-COR Biosciences #926-68071).

### *Trp53* status analysis

Genomic DNA from KPT tumors or tumor organoids were analysed in two independent PCR reactions. PCR A, adapted from Olive et al ^8^ enables detection of the GoF allele (330-bp band) and the Wt allele (290 bp band). PCR B allows detection of the LoF allele (370 bp band) and the second band (174 bp band) indicates presence of either Wt and/or GoF allele.

For both PCR’s, 100-150 ng of genomic DNA sample was used with PCR cycling conditions: 95 °C for 5 min, 35 cycles of 95 °C for 30 s, 55 °C (PCR A) or 53 °C (PCR B) for 30 s, and 72 °C for 30 s, final elongation step 72 °C for 5 min. Respective oligonucleotide primers (Supplementary Table 4) were used at a concentration of 500 nM in 2× PCR premix reagent (My Taq Red PCR Mix, Catalog #: MER-BIO-25044, Bioline).

### Kaplan-Meier survival analysis

All Kaplan–Meier survival analysis were performed with KMplot (KMplot.com) ^65^. RNA-seq data from stomach adenocarcinoma (N=375) patients within the pan-cancer dataset were analyzed. All other settings were kept as default.

### Bioinformatic analysis

#### Signaling pathway alteration analysis

Sanchez-Vega et al have analyzed all tumor types from the TCGA data set for frequency of signaling pathway alterations on a pathway level rather than individual gene mutations. Here, we have removed the esophageal tumor data from Sanchez-Vega et al’s Supplementary Table Files MMC1 and MMC4 ^27^. The resulting stomach adenocarcinoma selective data set was interrogated, and top alternated pathways are shown in Supplementary Fig. 1A.

#### Signaling pathway activation analysis

Signaling pathway activation analysis is based on RNA sequencing expression analysis of gene expression signatures. KRAS and PI3K pathway activation gene signatures were used as defined by Pek et al ^66^ and Zhang et al ^67^. TP53 pathway activation was not analyzed but TP53 genetic alteration frequencies are presented. Definition of a TP53 pathway activation gene signature is difficult, due to different gene signatures being associated with loss-of-function versus gain-of-function TP53 mutations as well as non-transcriptional activity of mutant TP53. TP53 mutation data were downloaded from the COSMIC database.

Signaling pathway activation analysis was adapted from Tan *et. al.* ^6^. Briefly, level 3 TCGA RNA-seq normalized data for 415 gastric cancer samples and 35 normal gastric samples, and their corresponding clinical information, were downloaded from the Broad Institute TCGA Genome Data Analysis Center Firehose. Gene expression values were log-transformed and centered to the standard deviation of the median across the samples included in the analysis. A μ score was calculated for each sample by averaging the standardized expression values of pathway signature genes. A pathway is deemed activated in a tumor sample if its μ score in that sample surpasses the 90th quantile of μ scores calculated across all normal samples.

#### IL6/Il11/GP130/STAT3 activation gene signature

STAT3 signaling can be induced by different stimuli and may result in activation of gene expression in different target gene subsets in a context dependent manner. Here we defined an IL6 and IL11 cytokine and GP130 dependent STAT3 signaling activation gene signature for gastric cancer. Previously we stimulated *Gp130^F/F^* mutant mice with either recombinant IL6 and IL11 and performed RNA sequencing and STAT3-Chip/Seq analysis ^31^. Here, we selected the top 12 upregulated genes from the IL11 stimulation and retained all genes which are equally upregulated in IL6 stimulated tumors and where STAT3 did bind to the gene body or promoter in the STAT3-ChipSeq analysis. Two genes were removed due to lack of human homologue genes. Four genes were removed due extensive non-tumor cell expression based on mouse stomach and *gp130^F/F^* tumor single cell RNA sequencing data ^68^ as well as human protein atlas expression distribution ^69^. One gene was removed due higher expression in normal tissue than STAD, resulting in a final five gene signature: *SOCS3*, *PIM3*, *OSMR*, *CLDN12* and *STAT3*.

### Statistical analysis

All experiments were conducted at least twice, if not otherwise indicated and for animal experiments with ≥3 sex-and aged-matched mice per group. For drug treatment experiments animals were randomized into treatment groups. Tumor growth measurements were performed blinded to treatment or genetic cohort conditions. No data was excluded from the analysis, if not indicated otherwise. Data used to generate the figure is provided in a Source File. GraphPad Prism 9 software was used to calculate means, standard error of the mean and was used to perform statistical testing. For two group comparisons, unpaired two-sided Student’s *t*-test was performed, either with or without Welch correction depending on deviation F of the data. If not normally distributed, Mann Whitney test was performed. Data comparison of more than two groups was done with one-way ANOVA, with multiple comparison testing by Tukey (when comparing the mean of each column with the mean of every other column), Dunnett (when comparing each column mean with the mean of a control column) or Sidak (when comparing the means of preselected pairs of columns). Kruskal-Wallis with Dunn’s multiple comparison testing was used for non-normally distributed data sets. Kaplan-Meier survival analysis Hazard ratios and p values were calculated with Mantel-Cox’s Log-rank test and contingency analyses was done using two-tailed Fisher’s exact test.

## Data availability

All relevant data are available from the corresponding author upon reasonable request.

## Acknowledgements

We thank David Lau and collaborators for the generation of the human GC TMAs. We thank the animal facility staff of La Trobe University and Austin Health for the care for our animals as well as the Austin Pathology department for histological support. We also acknowledge the support in flow cytometric cells sorting by David Baloyan (FACS facility, ONJCRI) and image acquisition and analysis by Sarah Ellis (Imaging Facility ONJCRI). Schematics for Figures 3B and Supplementary Fig. 7 were created with Biorender.com. Biorender publication licenses are provided and made out to our Cancer and Inflammation Program.

This work was supported through the National Health and Medical Research Council (NHMRC) Synergy Grant 2027459 (M.E), NHMRC Investigator grant 1173814 (M.E.), NHMRC Ideas grant 2020316 (M.B.), Victorian Cancer Agency midcareer Research Fellowship MCRF20018 (M.F.E.), AACR-Debbie’s Dream foundation Gastric Cancer Career Advancement Award 22-20-41-EISS (M.F.E.), La Trobe University Graduate Research Scholarship (A.H.). E.B. acknowledges support from ERC (ERC AdvG 884623) and AGAUR-2021-SGR-1278. We acknowledge support of the Victorian State Government Operational Infrastructure Support to the Olivia Newton-John Cancer Research Institute.

## Authors Contributions

Experimental Work: M.F.E, A.H., A.A., C.D., S.T., J.H., A.R.P., J.K., S.J., R.B., M.G.A, S.F., D.V.F.T., D.S.W., M.B.

Data Interpretation: M.F.E, A.H., A.A., C.D., S.T., R.B., Y.L., D.C., W.S., D.T., E.B., A.B., D.S.W., M.B., M.E.

Writing of Manuscript: M.F.E, A.H., A.A., W.S., E.B., D.S.W., M.B., M.E.

Study Concept: M.E., M.B., M.F.E

Funding: M.E., M.F.E, A.H., M.B.

