## Supplementary Information for "Mutant TP53 switches therapeutic vulnerability during gastric cancer progression within Interleukin-6 family cytokines"

**This PDF file includes:**

Supplementary Figures 1 to 7

Supplementary Tables 1 to 4

A

| Pathway | % mutated |
|---|---|
| Cell cyle | 65.8 |
| RTK/RAS | 59.5 |
| TP53 | 56.9 |
| PI3K | 41.5 |
| NOTCH | 39.2 |
| WNT | 37.3 |
| TGF-b | 28.7 |
| HIPPO | 22.2 |
| MYC | 18 |
| NRF2 | 1.6 |

B

| Pathway | % mutated |
|---|---|
| RTK/RAS | 59.5 |
| PI3K | 41.5 |
| TP53 | 56.9 |
| RTK/RAS + PI3K | 26.6 |
| RTK/RAS + TP53 | 36 |
| PI3K + TP53 | 24.5 |
| RTK/RAS + PI3K + TP53 | 15.1 |

C

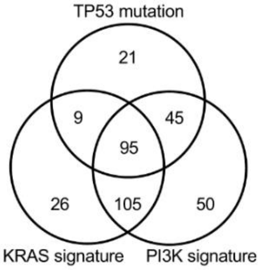


D

| Pathway | % activated |
|---|---|
| KRAS | 56.8 |
| PI3K | 71.2 |
| TP53* | 41 |
| KRAS + PI3K | 48.3 |
| KRAS + TP53* | 25.1 |
| PI3K + TP53* | 33.8 |
| KRAS + PI3K + TP53* | 23 |
| none | 15.2 |

* mutation

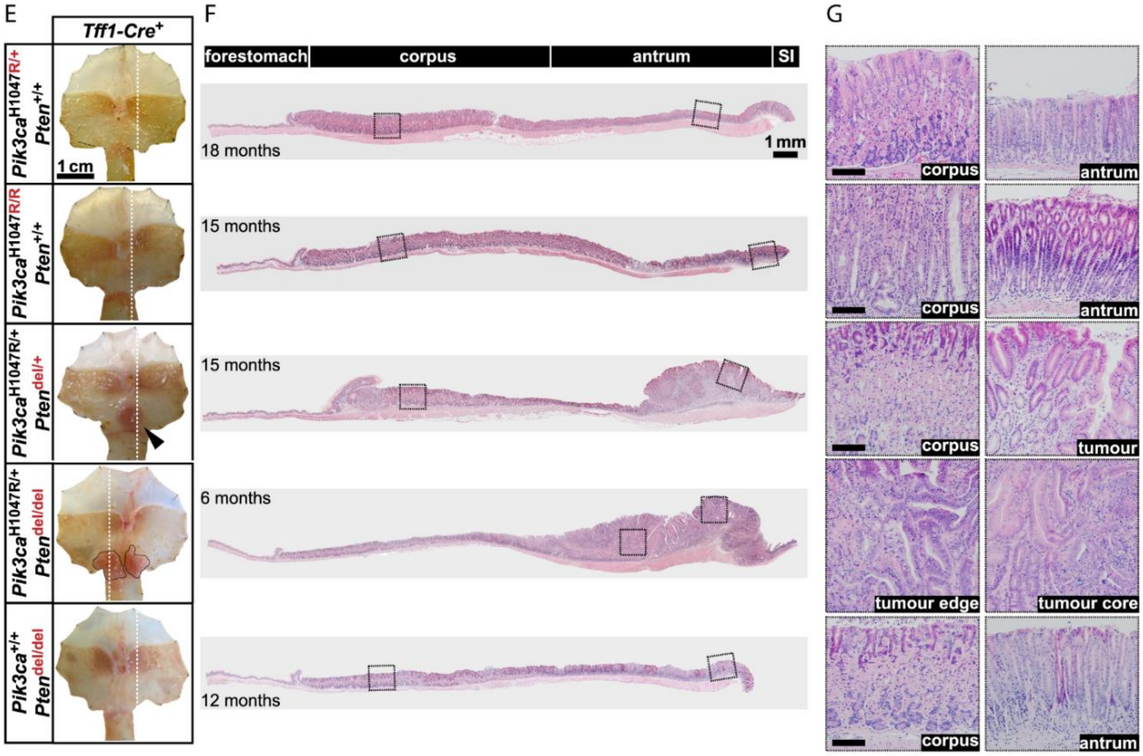


H

| Tff1-CreERT2-driven | Tumor (# GC/ # mice) | Carcinoma (# Carc/ # mice) | Metastasis (# metCarc/ # mice) |
|---|---|---|---|
| $Pik3ca^{H1047R/+};Pten^{+/+}$ | 0% (0/9) | 0% (0/9) | 0% (0/9) |
| $Pik3ca^{H1047R/H1047R};Pten^{+/+}$ | 0% (0/11) | 0% (0/11) | 0% (0/11) |
| $Pik3ca^{H1047R/+};Pten^{del/+}$ | 7.4% (2/27) | 0% (0/27) | 0% (0/27) |
| $Pik3ca^{H1047R/+};Pten^{del/del}$ | 93.8% (15/16) | 0% (0/16) | 0% (0/16) |
| $Pik3ca^{+/+};Pten^{del/del}$ | 0% (0/4) | 0% (0/4) | 0% (0/4) |

I

| Tff1-CreERT2-driven | Carcinoma (#Carc/# mice) | p value | Tis (# Tis/# Carc) | p value | T1 (# T1/# Carc) | p value | T2 (# T2/# Carc) | p value | T3 (# T3/# Carc) | p value | T4 (# T4/# Carc) | p value |
|---|---|---|---|---|---|---|---|---|---|---|---|---|
| $Kras^{G12D/+}$; $Pik3ca^{H1047R/+};Pten^{del/+}$ | 56% (9/16) | 0.76 | 22% (2/9) | 0.095 | 56% (5/9) | 0.001 | 11% (1/9) | 0.32 | 11% (1/9) | 1 | 0% 0/9) | 0.0001 |
| $Kras^{G12D/+}$; $Pik3ca^{H1047R/+};Trp53^{R172H/+}$ | 61.3% (19/31) | | 0% (0/19) | | 0% (0/19) | | 0% (0/19) | | 10.5% (2/19) | | 89.5% (17/19) | |

**Supplementary Figure 1. Frequency of KRAS, PI3K and TP53 pathway alterations in human gastric cancer and PI3k pathways potency as cancer driver in mice**

**A** Top 10 mutated (in percent) oncogenic pathways in stomach adenocarcinoma (STAD) patients. Data extracted from TCGA data set (STAD n=383). RTK/RAS= Receptor tyrosine kinase/RAS.

**B** Table shows selected pathways mutation frequencies (as in **A**) for single, double and triple pathway involvement.

**C** Venn diagram for oncogenic signaling pathway activation gene signature analysis, showing number of patients with STAD with KRAS and/or PI3K pathway activation (based on gene expression signature) and *TP53* gene mutation (silent mutation (n=1) excluded). 63 patients had neither *TP53* mutation nor KRAS or PI3K activation status tumors (STAD n=414).

**D** Data from **C** presented in table format, showing the precent frequency of patients with single, double, and triple pathways activation (*TP53 data is gene mutation based).

**E** Representative whole-mount stomachs from Cre-positive mice of the indicated genotypes, collected at 6-18 months (as indicated) after Tamoxifen administration (*arrowhead* and *black lines* indicate tumors).

**F,G** H&E-stained cross-sections, cut along the dotted line (in **D**), extend from the forestomach to the proximal end of the small intestine (SI) (**E**). Boxed regions are shown in higher magnifications (**F**). Scale bars 100µm.

**H** Frequency (in percent) of tumor development in Cre-positive mice of the indicated genotype.

**I** Table shows invasive stages (T stage) of $Kras^{LSL\text{-}G12D/+}$; $Pik3ca^{LSL\text{-}H1047R/+}$;$Pten^{flox/+}$ and $Kras^{LSL\text{-}G12D/+}$; $Pik3ca^{LSL\text{-}H1047R/+}$;$Trp53^{LSL\text{-}R172H/+}$ (*KPT*) carcinoma bearing mice. Tis = carcinoma in situ. P values of Fisher's exact test are shown.

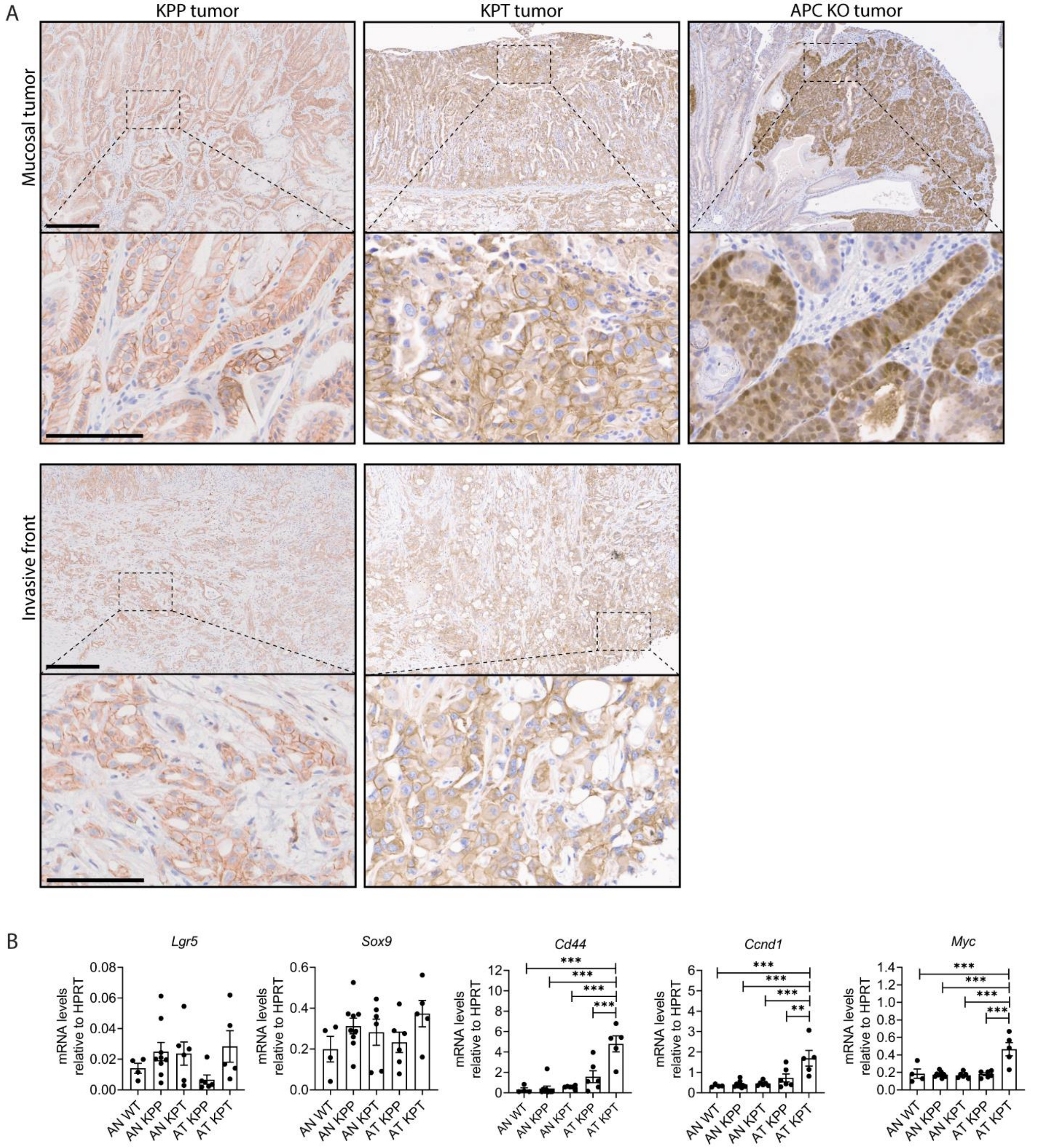


**Supplementary Figure 2. Nuclear β-Catenin and WNT-gene signature analysis in tumors without overt canonical WNT signaling pathway mutations**

**A** Representative microscopy images of anti-β–catenin IHC stained tumors of *KPP*, *KPT* and $Apc^{KO}$ ($Tff1^{CreERT2};Apc^{fl/fl}$) mice. *KPP* and *KPT* image represent strongest staining (of n=10 mice analyzed). $\text{Apc}^{\text{KO}}$ shown as positive control for nuclear β–catenin positivity; $Apc^{KO}$ mice develop non-invasive adenomas. Scale bars equal 300 µm (top images) and 100 µm (bottom images).

**B** qPCR analysis of transcription levels of canonical WNT signaling target genes of whole tumor tissues. Stomach antrum normal (AN) and antrum tumors (AT) of indicates mutant mice were analyzed. Mean±SEM. One-way ANOVA with Tukey's multiple comparisons test was performed; ns=non-significant, * =  p<0.01, ** = p<0.001, *** = p<0.0001. Each data points represents one mouse; N = 4, 9, 6, 6 and 5 (for each graph).

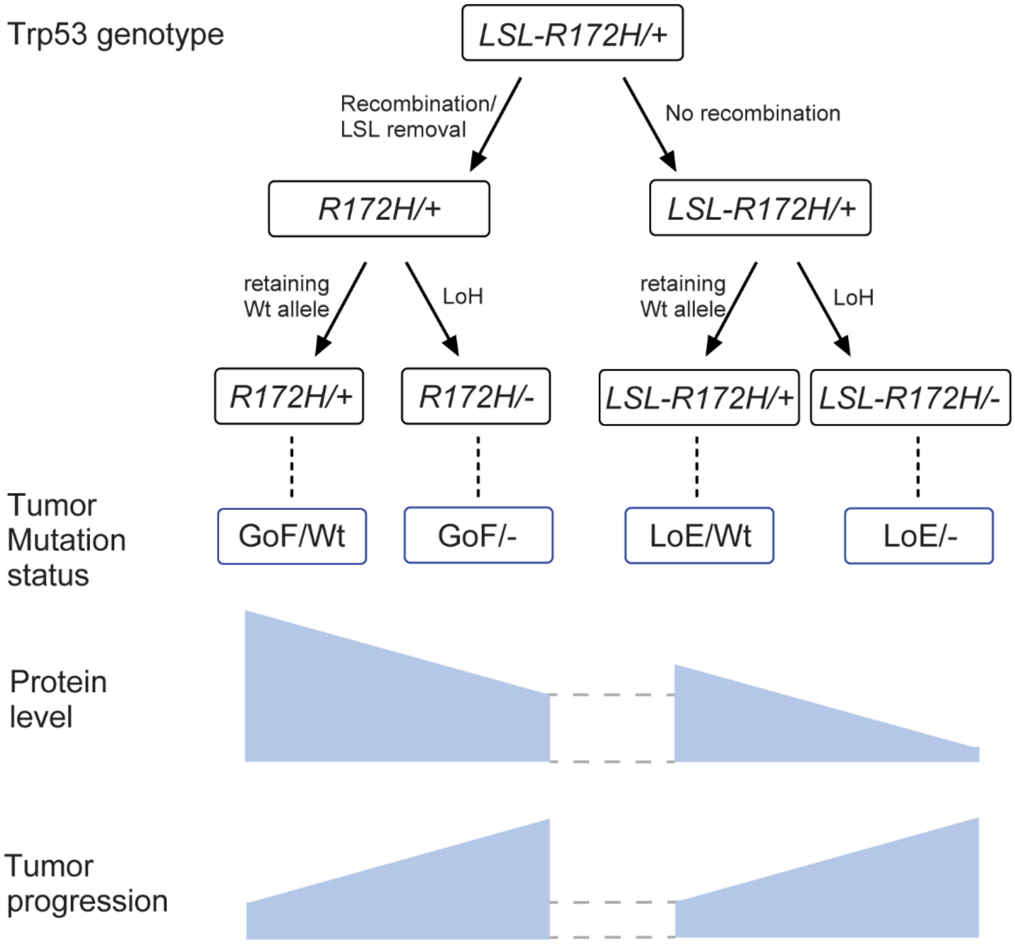


**Supplementary Figure 3. TP53 mutations status at genetic, protein and phenotype level**

Upon tamoxifen administration Tff1$^{CreERT2}$ recombinase drives removal of the Los-Stop-Lox (LSL) cassette from the LSL-R172H allele to yield the R172H mutant expression, which is classified a Gain-of-Function (GoF) mutant. Some cells do not undergo Cre-recombination and retain the unrecombined LSL-R172H allele, which is a Loss-of-Expression (LoE) mutant (or "null' allele) as no or minimal TP53 protein can be expressed from this allele. Throughout tumorigenesis and/or tumor progression the wildtype (Wt) Trp53 allele can be genetically lost (-), also called Loss-of-heterozygosity.

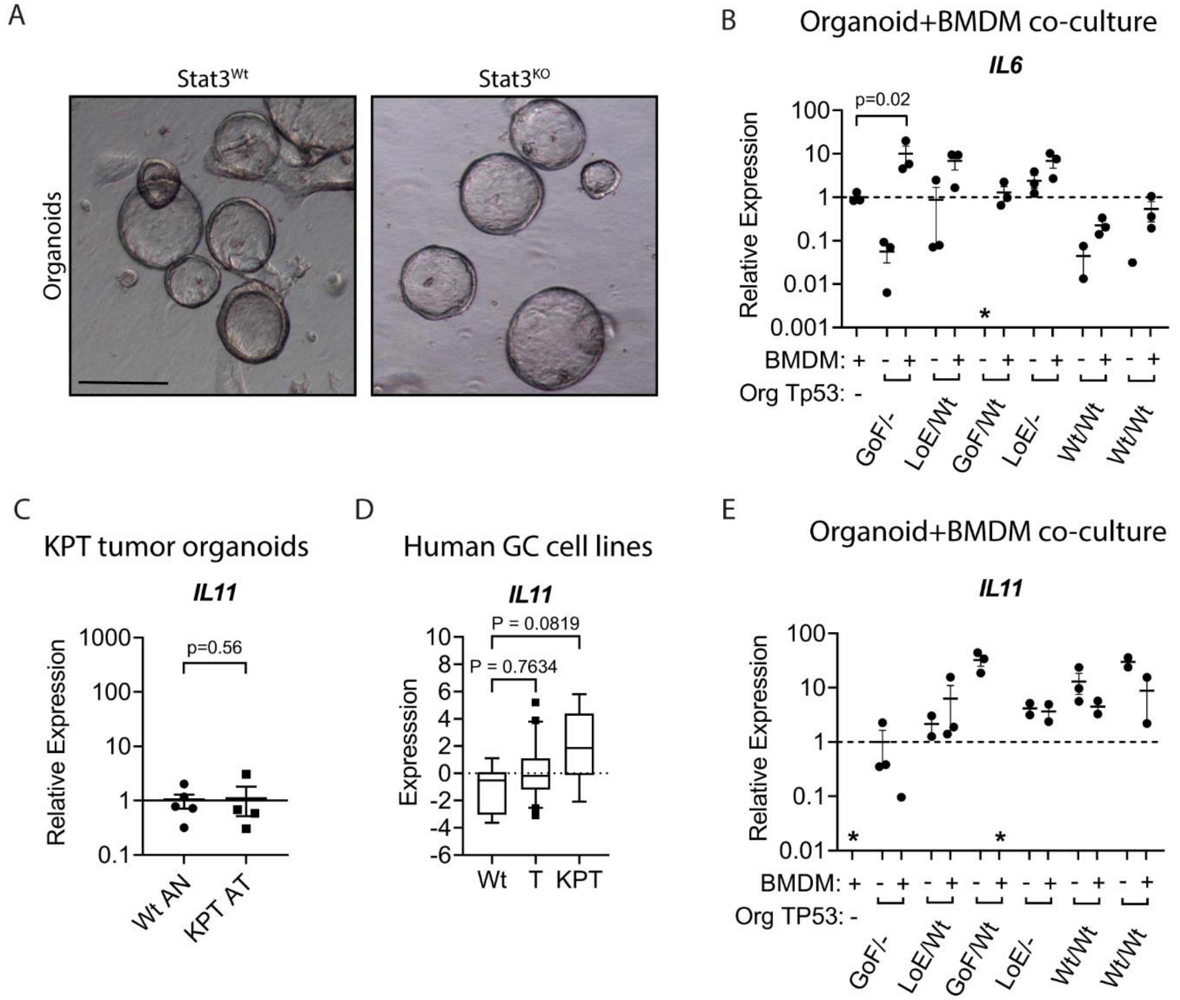


**Supplementary Figure 4. IL6 and IL11 expression profile in mouse KPT tumor organoids and human gastric cancer cells**

**A** Representative brightfield microscopy images of *Stat3*$^{Wt}$ and *Stat3*$^{KO}$ GC organoids (scale bar = 200µm).

**B** qPCR analysis for *IL6* expression of either bone marrow derived macrophages (BMDM) alone, tumor organoids alone (with indicated *TP53* status) or BMDM + organoids co-cultured. The first four organoid cultures are derived from tumors of *KPT* mice and the two *Trp53*$^{wt/wt}$ organoids are derived from *gp130*$^{Y757F/Y757F}$ mouse tumors. Each data point presents the mean of technically triplicates from three independent experiments (some *Trp53*$^{WT/WT}$ organoids alone samples had *IL6* expression below detection limit). Relative expression normalized to BMDM alone. Mean±SEM. One-way ANOVA with Dunnett's multiple comparisons test calculated p value shown, all other comparisons resulted in p>0.2.

**C** qPCR-determined expression levels of *IL11* in organoids derived from wild type antrum stomach (Wt) and stomach tumors of *KPT* mutant mice. Each data points represents organoid derived from different mice. P-values of Mann Whitney test are shown. N = 4 and 4.

**D** *IL11* expression levels in human gastric cancer cell lines grouped into *TP53*$^{wt}$ (*Wt*), *TP53* (*T*) mutant or *KRAS;PI3K;TP53* (*KPT*) mutant activation signature positive. RNA sequencing data was downloaded from the Broad Institutes Cancer Cell Line Encyclopedia. Data is shown as Box & Whiskers plots (10-90 percentile), comprising WT n=8, T n=21 and KPT n=7 data points. Kruskal-Wallis test and Dunn's multiple comparison's test were performed.

**E** *IL11* expression analysis from BMDM + organoid co-culture experiment as in **B**. Asterisk (*) indicates sample groups were raw values of each technical replicates from three independent experiments were below the detection limit! Relative expression normalized to GoF/- organoid alone, as *IL11* in BMDM alone expression was below detection limit. Data presentation and statistical testing as in **B**.

A

| Tff1-CreRT2-driven | Carcinoma (#Carc/# mice) | p value | Tis (# Tis/# Carc) | p value | T1 (# T1/# Carc) | p value | T2 (# T2/# Carc) | p value | T3 (# T3/# Carc) | p value | T4 (# T4/# Carc) | p value |
|---|---|---|---|---|---|---|---|---|---|---|---|---|
| $Kras^{G12D/+}$; $Pik3ca^{H1047R/+}$;$Trp53^{R172H/+}$ | 61.3% (19/31) | 0.77 | 0% (0/19) | | 0% (0/19) | | 0% (0/19) | | 10.5% (2/19) | 0.63 | 89.5% (17/19) | 0.63 |
| $Kras^{G12D/+}$; $Pik3ca^{H1047R/+}$;$Trp53^{R172H/+}$;$Il11ra^{+/-}$ | 66.7% (12/18) | | 0% (0/12) | | 0% (0/12) | | 0% (0/12) | | 16.7% (2/12) | | 83.3% (10/12) | |

B

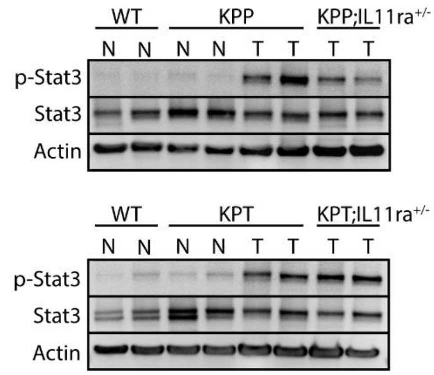


C

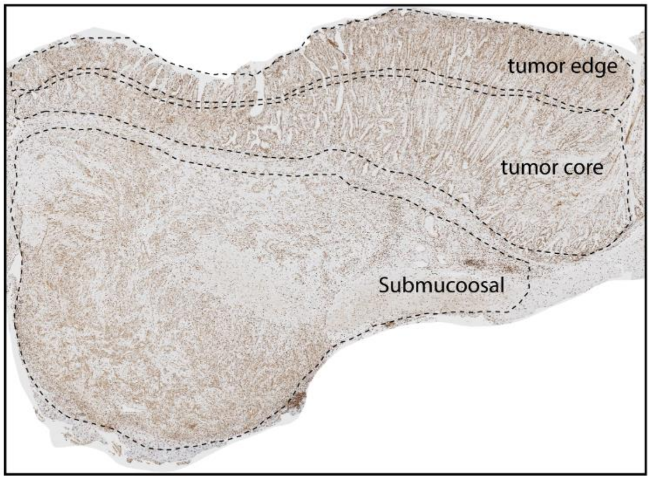


D

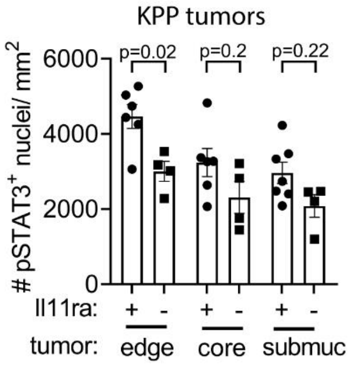


E

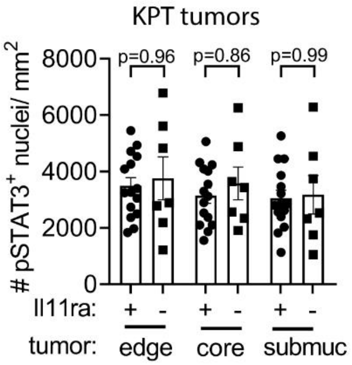


**Supplementary Figure 5. Mutant *Trp53* renders Stat3 signaling activity IL11 independent in *KPT* mutant tumors**

**A** T stage table illustrates depth of invasion for carcinomas with $Kras^{LSL\text{-}G12D/+}$; $Pik3ca^{LSL\text{-}H1047R/+}$;$Trp53^{LSL\text{-}R172H/+}$ (*KPT*) versus $Kras^{LSL\text{-}G12D/+}$; $Pik3ca^{LSL\text{-}H1047R/+}$;$Trp53^{LSL\text{-}R172H/+}$;$Il11ra^{+/-}$ genotypes. Tis = carcinoma in situ. P values of Fisher's exact test are shown.

**B** Immunoblotting for pSTAT3 (pY757), total-STAT3 and Actin from tissue lysates from antrum normal (*N*) or tumor (*T*) of wild type (*WT*), *KPP* and *KPP;IL11ra*$^{+/-}$ mice (upper plots) and wild type (*WT*), *KPT* and *KPT;IL11ra*$^{+/-}$ mice (lower plots).

**C** Representative image of pSTAT3 immunohistochemical stained *KPP* tumor (same as in Fig. 3A) exemplifying the definition of tumor edge, tumor core and submucosal stomach layers containing invaded tumor cells.

**D, E** Quantification of immunohistochemical staining for p-Stat3 in FFPE-sections of *KPP* tumors (C) and *KPT* tumors (D). Analyzed tumors were either IL11ra$^{+/+}$ (+) or IL11ra$^{+/-}$ (-). Quantification was performed in tumor edge, tumor core and tumor invaded submucosal layers (as outlined in **B**). P values of one-way ANOVA with Sidak's multiple comparison test are shown. Each data point represents an individual mouse. N = 6, 4 (**D**) and 15, 7 (**E**)(for each edge, core and submuc).

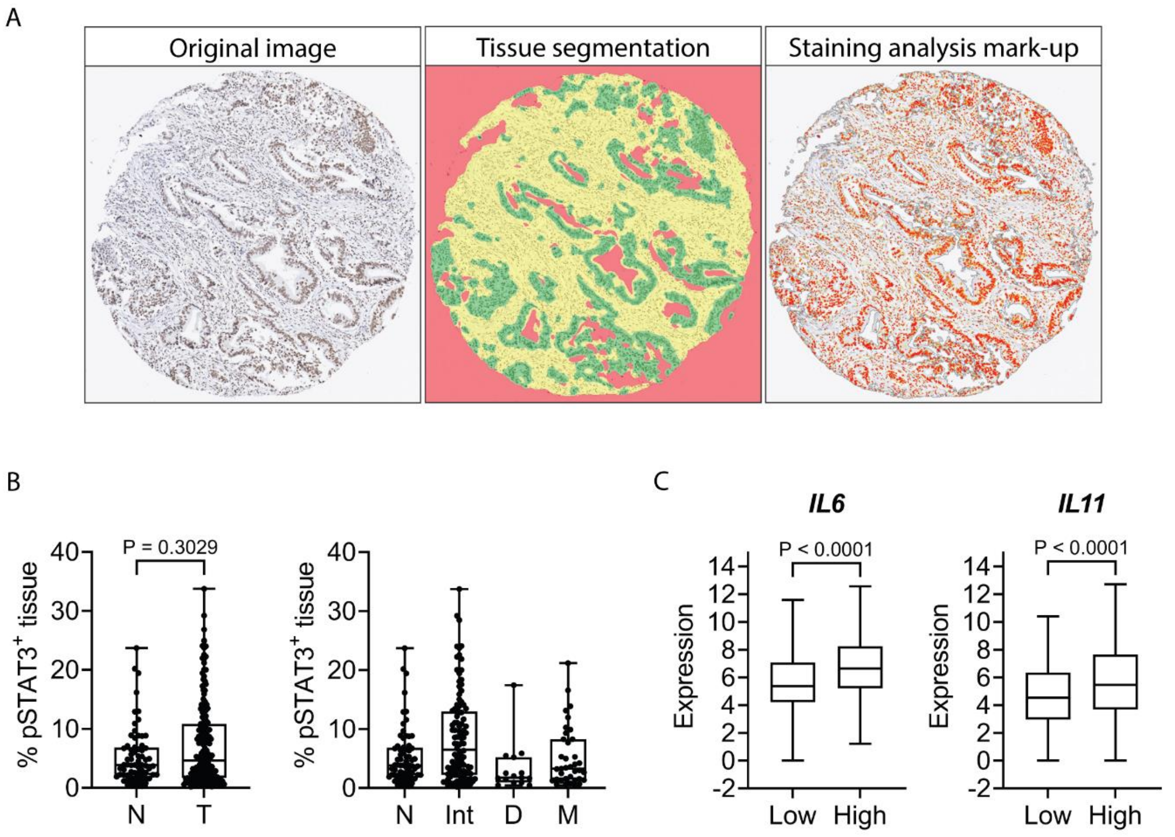


**Supplementary Figure 6. STAT3 signaling quantification and *IL6* cytokine expression on human gastric cancer Tissue Micro Arrays**

**A** Halo analysis of pSTAT3 stained human GC Tumor Micro array (TMA). Left image is the original pSTAT3 image of one representative tumor core of the TMA. Middle image shows the tissue segmentation: halo AI classified glass/no tissue (red), tumor stroma (yellow) and tumor (green). Right image presents Halo staining quantification Make-up, with weak (yellow), intermediate (orange) and strong (red) pSTAT3 positivity.

**B** Halo AI staining quantification of pSTAT3 on total tumor core of human GC TMAs (without Halo tissue segmentation). Left graph Box plots: Min to Max showing all data points. N= normal stomach, T= Tumor, Int=Intestinal GC, D=diffuse GC, M=Mixed GC. left: Mann-Whitney, right: Kruskali-Wallis+Dunnet's multiple Comparisons test (all p>0.11). N = 68, 168 (left) and 68, 102, 15 and 34 (right).

**C** mRNA Expression of IL6 and IL11 in Stat3 activation gene signature negative versus positive stomach adenocarcinoma samples (from TCGA data set). Boxplot with error bars showing min to max, p values shown were calculated with Mann-Whitney test. N = 280, 167 (both graphs).

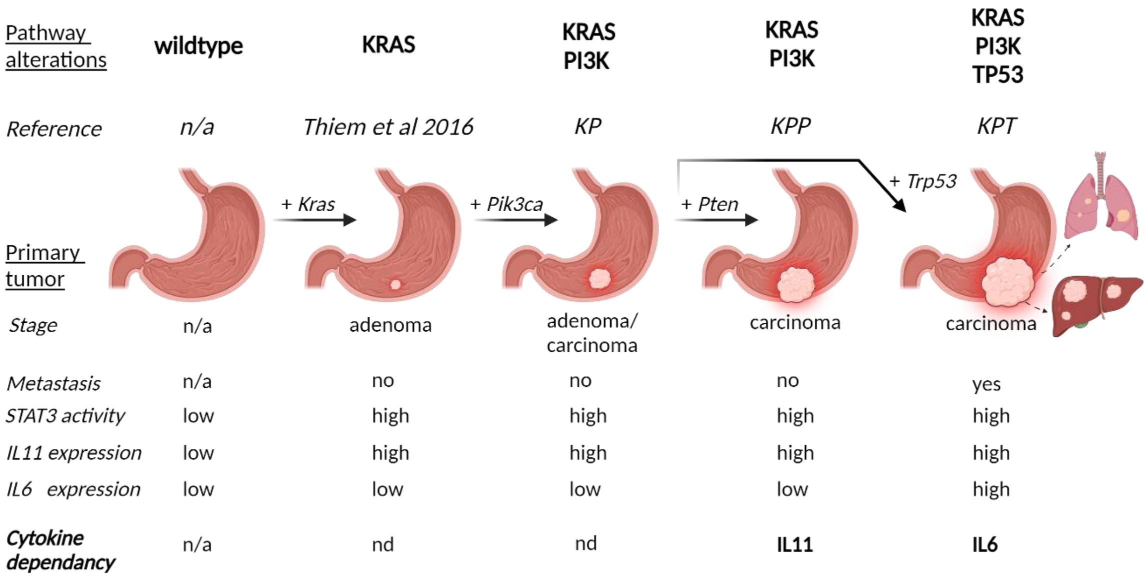


**Supplementary Figure 7. Graphical summary of TP53 mutation driven IL-6 family cytokine dependency switch**

Illustrates the pathway activation series with corresponding gastric tumor progression, associated *IL11*, *IL6* expression, STAT3 signalling activity and resulting cytokine dependencies. n/a = not applicable, nd = not determined. This figure was created with Biorender.com.

**Supplementary Table 1. Histopathological assessment of stomach HE's of tumor-bearing KPT mice**

| Diagnosis | WHO subtype | Lauren's subtype | Differentiation grade | Invasion (on slide) | Additional observations |
|---|---|---|---|---|---|
| **STAD** | tubular | intestinal | poor | serosa | |
| **STAD** | tubular | intestinal | moderate/poor | serosa | |
| **STAD** | mixed | intestinal | poor | subserosa | |
| **STAD** | solid | intestinal | poor | serosa | spans GOJ |
| **STAD** | solid | intestinal | poor | serosa | lymph node metastases |
| **STAD** | mixed | intestinal | moderate/poor | invades liver directly | spans GOJ |
| **STAD** | solid | intestinal | poor | serosa | |
| **STAD** | solid | intestinal | poor | serosa | |
| **STAD** | tubular | intestinal | poor | invades liver directly | lymph node metastasis |
| **STAD** | tubular | intestinal | moderate | subserosa | lymph node metastasis |
| **STAD** | tubular | intestinal | poor | into muscularis propria | |
| **STAD** | tubular | intestinal | poor | serosa | |
| **STAD** | tubular | intestinal | poor | subserosa | |
| **STAD** | tubular | intestinal | poor | invades pancreas directly | |
| **STAD** | tubular | intestinal | moderate | subserosa | lymph node metastasis |
| **STAD** | tubular | intestinal | poor | invades pancreas directly | |
| **STAD** | tubular | intestinal | poor | serosa | |
| **STAD** | tubular | intestinal | moderate | serosa | |
| **STAD** | tubular | intestinal | poor | invades pancreas directly | |

STAD = stomach adenocarcinoma, mixed = mixed tubular and solid WHO subtype, GOJ = Gastric-oesophageal junction

**Supplementary Table 2. Clinicopathological characteristics of GC patient stratified by tumor-compartment P-STAT3 IHC staining.**

| **Characteristic** | **$pSTAT3^{Low}$ N= 84** | **$pSTAT3^{High}$ N= 84** | **p value** |
|---|---|---|---|
| **Age (years)** | | | |
| Mean ± SD | 68.31 ± 10.67 | 70.31 ± 11.91 | 0.1 |
| Median | 70 | 72 | |
| Range | 44 - 92 | 27 – 87 | |
| **Site** | | | |
| Gastric | 68 (81) | 63 (75) | 0.46 |
| GOJ | 16 (20) | 21 (25) | |
| **Lauren subtype** | | | |
| Intestinal | 43 (57) | 59 (79) | 0.013 |
| Diffuse | 11 (15) | 4 (5) | |
| Mixed | 22 (29) | 12 (16) | |
| **AJCC TNM stage** | | | |
| 1 | 9 (11) | 13 (16) | 0.13 |
| 2 | 24 (30) | 34 (42) | |
| 3 | 40 (51) | 26 (32) | |
| 4 | 6 (8) | 8 (10) | |
| **Grade** | | | |
| Low | 17 (20) | 30 (36) | 0.03 |
| High | 67 (80) | 53 (64) | |
| **Lymph node involvement** | | | |
| No | 24 (29) | 36 (43) | 0.08 |
| Yes | 58 (71) | 47 (57) | |
| **T4 stage** | | | |
| No | 50 (61) | 61 (73) | 0.1 |
| Yes | 32 (39) | 22 (27) | |
| **DNA Mismatch Repair status** | | | |
| Proficient | 66 (80) | 69 (82) | 0.7 |
| Deficient | 17 (20) | 15 (18) | |

P values are shown from Mann Whitney (Age), Chi-square (AJCC stage, Lauren subtype), Fisher exact (all remaining). Column percentages are shown in brackets. GOJ = Gastric Oesophageal Junction, AJCC= American Joint Committee on Cancer.

**Supplementary Table 3. Oligonucleotide sequences for quantitative RT-PCR**

| Gene | Forward Primer | Reverse Primer |
|---|---|---|
| *Il6* | TCTATACCACTTCACAAGTCGGA | GAATTGCCATTGCACAACTCTTT |
| *Il11* | ACT TCC ACC TCA GGA CAT CG | GGT GGT TAA AAG GAC AGG CA |
| *Gapdh* | CAC TGA GCA TCT CCC TCA CA | GTG GGT GCA GCG AAC TTT AT |
| *Lgr5* | CCT TGG CCC TGA ACA AAA TA | ATT TCT TTC CCA GGG AGT GG |
| *Sox9* | CCA CGG AAC AGA CTC ACA TCT CTC | CTG CTC AGT TCA CCG ATG TCC ACG |
| *Cd44* | GTC TGC ATC GCG GTC AAT AG | GGT CTC TGA TGG TTC CTT GTT |
| *Ccnd1* | TCG TGG CCT CTA AGA TGA AGG A | TCG GGC CGG ATA GAG TTG T |
| *Myc* | AGC TGT TTG AAG GCT GGA TT | AAT AGG GCT GTA CGG AGT CG |
| *Hprt* | GGC CAG ACT TTG TTG GAT TTG | CGC TCA TCT TAG GCT TTG TAT TTG |

**Supplementary Table 4. Primer sequences used for PCR A and B for Trp53 status analysis**

| Primer Name | Primer Sequence |
|---|---|
| **PCR A** | |
| Olive2004_LSLcas-Recomb_F | agc ctg cct agc ttc ctc agg |
| Olive2004_LSLcas-Recomb_R | ctt gga gac ata gcc aca ctg |
| **PCR B** | |
| Trp53 WT | agg tgt ggc ttc tgg ctt c |
| Trp53 neo mutant | cca tgg ctt gag taa gtc tgc a |
| Trp53 common | gaa act ttt cac aag aac cag atc a |
